# PITHD1: An Endogenous Inhibitor of the 26S Proteasome During Cellular Dormancy

**DOI:** 10.1101/2024.12.04.626795

**Authors:** Sascha J. Amann, Ken Dong, Josef Roehsner, Dominik Krall, Irina Grishkovskaya, Harald Kotisch, Alexander Schleiffer, Elisabeth Roitinger, Andrea Pauli, Andreas Martin, David Haselbach

## Abstract

Cellular dormancy represents a state of regulated growth arrest essential for diverse biological processes, from reproduction to cancer progression. While mechanisms controlling protein synthesis in dormant cells have been identified, how cells regulate protein degradation during dormancy remains unclear. Using zebrafish oocytes, eggs and embryos as a model system, we discovered PITHD1 as an endogenous inhibitor of the 26S proteasome. Our high-resolution cryoEM structure reveals that PITHD1 simultaneously blocks three crucial functional sites on the 19S regulatory particle which are required for ubiquitin recognition, processing, and substrate translocation. This triple-lock mechanism effectively prevents protein degradation in dormant cells. Given PITHD1’s evolutionary conservation across species, this mechanism likely represents a general strategy for reversible proteasome regulation during cellular dormancy. Our findings establish a new paradigm for controlling proteostasis in quiescent states.

---

Cellular dormancy occurs when cells exit the active cell cycle and enter a state of growth arrest and reduced metabolic activity. This cell state ensures the conservation of energy and resources until cells receive a stimulus to re-enter the cell cycle. As such, cellular dormancy plays a critical role in a wide range of biological processes, such as tissue homeostasis, repair and regeneration, immune response, ageing and reproduction (*1*). Furthermore, cellular dormancy, also referred to as senescence, has important implications in cancer, where cancer cell dormancy has been linked to resistance to treatment and disease recurrence (*2*).

In general, entering a dormant state involves a variety of molecular changes, such as changes in gene expression, epigenetic regulation, metabolic activity, energy homeostasis and proteostasis (*3*). Over the years, most studies have focused on understanding changes in signaling pathways, metabolism and gene expression that accompany dormancy; however, mechanisms that dormant cells use to maintain their protein composition over extended periods remain unclear.

In most cases, the levels of a given protein reflect the balance between ribosome-mediated synthesis and proteasome-mediated degradation. In the context of cellular dormancy, we recently showed that dormant eggs feature translationally inactive ribosomes that are inhibited through association with a network of “dormancy factors” (*4*). Furthermore, a recent study of dormant mammalian oocytes revealed that extreme protein stability is involved in maintaining the female germline (*5*), indicating that a dormant state is associated with unique proteostasis requirements. In agreement with this general insight, cytoplasmic proteasomes in quiescent immature oocytes were observed to be stored in endolysosomal vesicular assemblies and kept inactive (*6*). Collectively, these findings suggest that both protein synthesis and protein degradation need to be tightly regulated in quiescent states. However, the mechanisms underlying this modulation remain poorly understood.

Here, we identified PITHD1 as a protein that is associated with the 26S proteasome in dormant zebrafish oocytes and eggs. Our structural and functional analysis revealed that PITHD1 inactivates the proteasome by engaging with the 19S regulatory particle and blocking ubiquitin recognition, processing, and substrate translocation. We also identified a possible tyrosine phosphorylation-dependent mechanism for PITHD1 regulation, that may serve to restore proteasomal protein degradation upon exit from dormancy. While PI31 has been described as an inhibitor of the 20S Core Particle (CP) (*7*), PITHD1 is the first identified endogenous inhibitor of the 19S Regulatory Particle (RP) of the 26S proteasome.

## Identification of PITHD1 as proteasomal dormancy factor

Gene expression is repressed in mature zebrafish eggs and only activated after fertilisation and resumption of cell proliferation in embryos (*4*). Many factors involved in regulating the egg-to-embryo transition in zebrafish are evolutionarily conserved, making zebrafish an appropriate model to study these processes. To investigate proteasomes during oogenesis and embryonic development, we isolated proteasomes from zebrafish ovaries, eggs and embryos via sucrose gradient centrifugation of cell extracts (Fig. 1A). Gradient fractions containing proteasomes were subjected to chymotrypsin-like activity assays with normalised proteasome levels. We observed that proteasomal activity was decreased in ovaries and eggs compared to rapidly dividing embryos (Fig. 1B, fig S1A, B), suggesting that the activity of proteasomes pre-fertilisation may be inhibited by the association with “dormancy factors”.

**Fig. 1.**
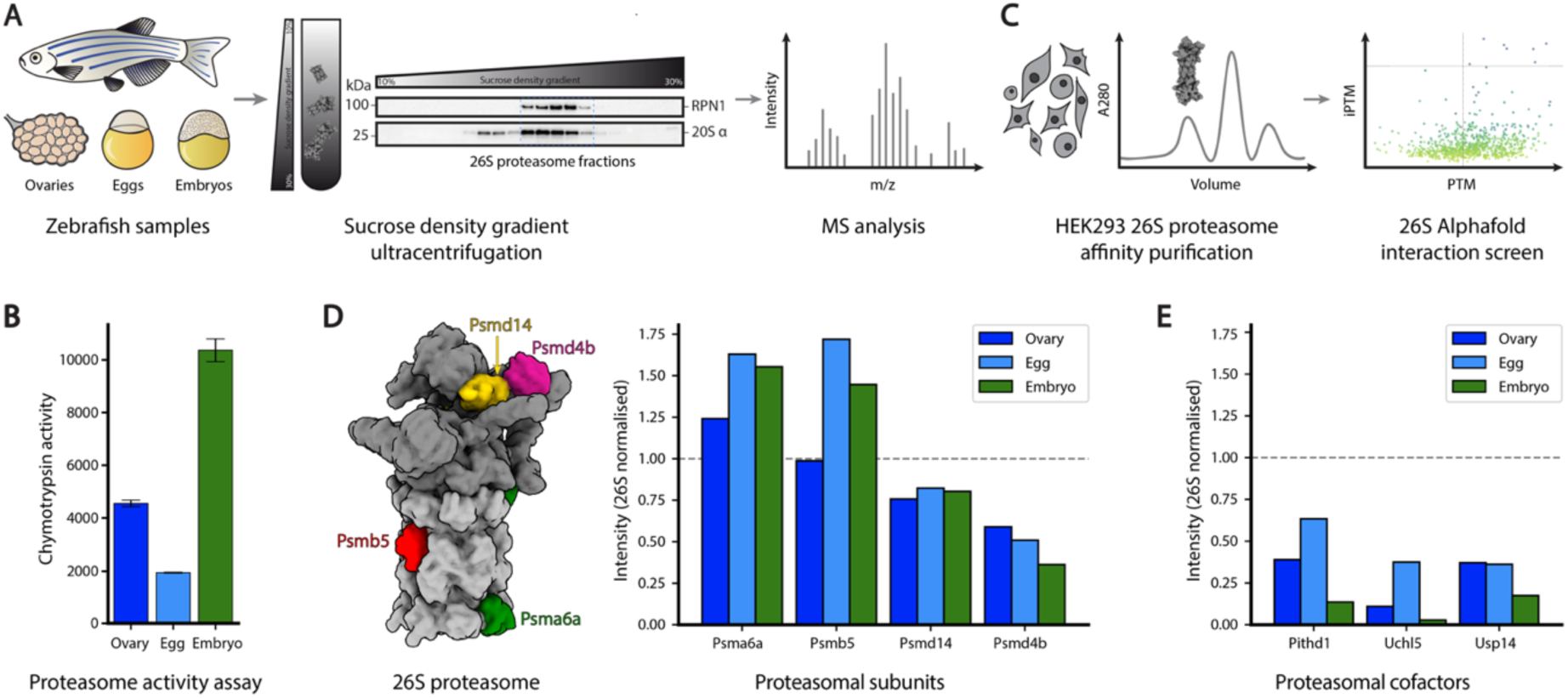
Identification of Pithd1 as a proteasome dormancy factor. (A) Extracts from three stages of zebrafish development were lysed and proteasomes were enriched by sucrose density gradient ultracentrifugation. (B) 26S proteasome fractions were tested for proteasome activity using a fluorogenic substrate (n = 3). (**C**) AlphaFold interaction screen against the 26S proteasome using proteins pulled down by HEK293 26S proteasome affinity purification to identify potential interaction partners. (**D**) 26S proteasome with the 20S CP in light grey with highlighted subunits Psma6a (green), Psmb5 (red) and the 19S RP in dark grey with highlighted subunits Psmd4b (purple), Psmd14 (yellow) and their corresponding Mass-spectrometry intensities in the zebrafish sample 26S proteasome fractions. (**E**) Mass spectrometry intensities of the most abundant potential 26S proteasomal cofactors in the zebrafish sample 26S proteasome fractions identified in (**C**).

To identify potential dormancy factors of the proteasome, we performed a quantitative proteomic analysis of these proteasomes purified from each developmental stage. We cross-referenced these results with proteins identified from affinity-purified 26S proteasomes in HEK293 cells to focus on factors specifically associated with the proteasome. To predict potential proteasomal cofactors, we performed an AlphaFold2 screen against all constitutive proteasomal subunits, validated using AlphaFold3 multimer predictions (*8*, *9*) (Fig. 1C).

This analysis revealed several proteins that were either enriched or depleted in their association with proteasomes from pre-fertilisation samples. 8 proteins in ovaries and 13 proteins in eggs show at least a two-fold enrichment on proteasomes compared to embryos (Table S1), highlighting them as potential “dormancy factors”. Interestingly, Akirin2 was amongst the down-regulated potential proteasomal cofactors in ovaries and eggs (Table S2). In light of our previous work which revealed an important function for Akirin2 in nuclear import of proteasomes during cell cycle progression (*10*), the observed depletion of Akirin2 from proteasomes isolated from pre-fertilization samples indicates the quiescence of the ovary and egg samples. Three proteins, upregulated in the dormant pre-fertilization samples and present at the same order of magnitude as the 19S RP subunits, were predicted to interact specifically with the 19S regulatory particle: Pithd1, Uchl5, and Usp14 (Fig. 1D–E). AlphaFold3 multimer predictions further supported their interaction with functional sites of the 19S RP (fig. S1D) (*11*).

Both USP14 and UCHL5 are well-characterized proteasomal deubiquitinases (DUBs). While UBP14 can inhibit proteasomal activity and also provides an alternative deubiquitination pathway to RPN11 (*12*), UCHL5 prevents protein degradation by removing ubiquitin chains from recruited substrates (*13–15*). In contrast, although PITHD1 (Proteasome Interacting Thioredoxin Domain 1) remains functionally uncharacterized despite being highly conserved among vertebrates (fig. S2A), exemplifying a member of the “dark proteome.” Given PITHD1’s particular enrichment on proteasomes isolated from dormant zebrafish ovaries and eggs and depletion in embryos (Fig. 1E, Table S1), along with its predicted interactions with multiple regulatory sites of the proteasome (fig. S1D), we focused our subsequent analyses on PITHD1.

## Structure of PITHD1 bound to the 26S proteasome

PITHD1 is highly conserved as it is found in most eukaryotes and highly expressed in human oocytes (fig. S2A, (*16*)). To understand how PITHD1 regulates proteasome activity, we reconstituted the human 26S proteasome-PITHD1 complex and determined its structure using cryoEM. The 26S proteasome functions through an ordered sequence of conformational states that facilitate substrate processing. In its ground state (S1), the regulatory particle maintains a compact arrangement where the DUB RPN11 is offset from the central channel. Substrate binding favours a transition through an intermediate state (S2), marked by RPN1 rotation and initial substrate engagement. Subsequently, ATP-dependent conformational changes drive the proteasome into activated states (S3/S4), repositioning RPN11 above the ATPase channel to couple deubiquitination with substrate translocation. Understanding how PITHD1 affects these states was crucial for deciphering its inhibitory mechanism. Previous structural studies have shown that regulatory factors can stabilize specific states, thereby controlling proteasome function (*17*).

Reconstitution by sucrose gradient ultracentrifugation (fig. S2B) and an optimized data processing pipeline (*18*) yielded a high-resolution map (2.3 Å overall) of a PITHD1-proteasome complex in the S1 state, with the PITHD1-proteasome interface resolved at 2.5 Å (Fig. 2A-C, fig. S3, fig. S4).

**Fig. 2.**
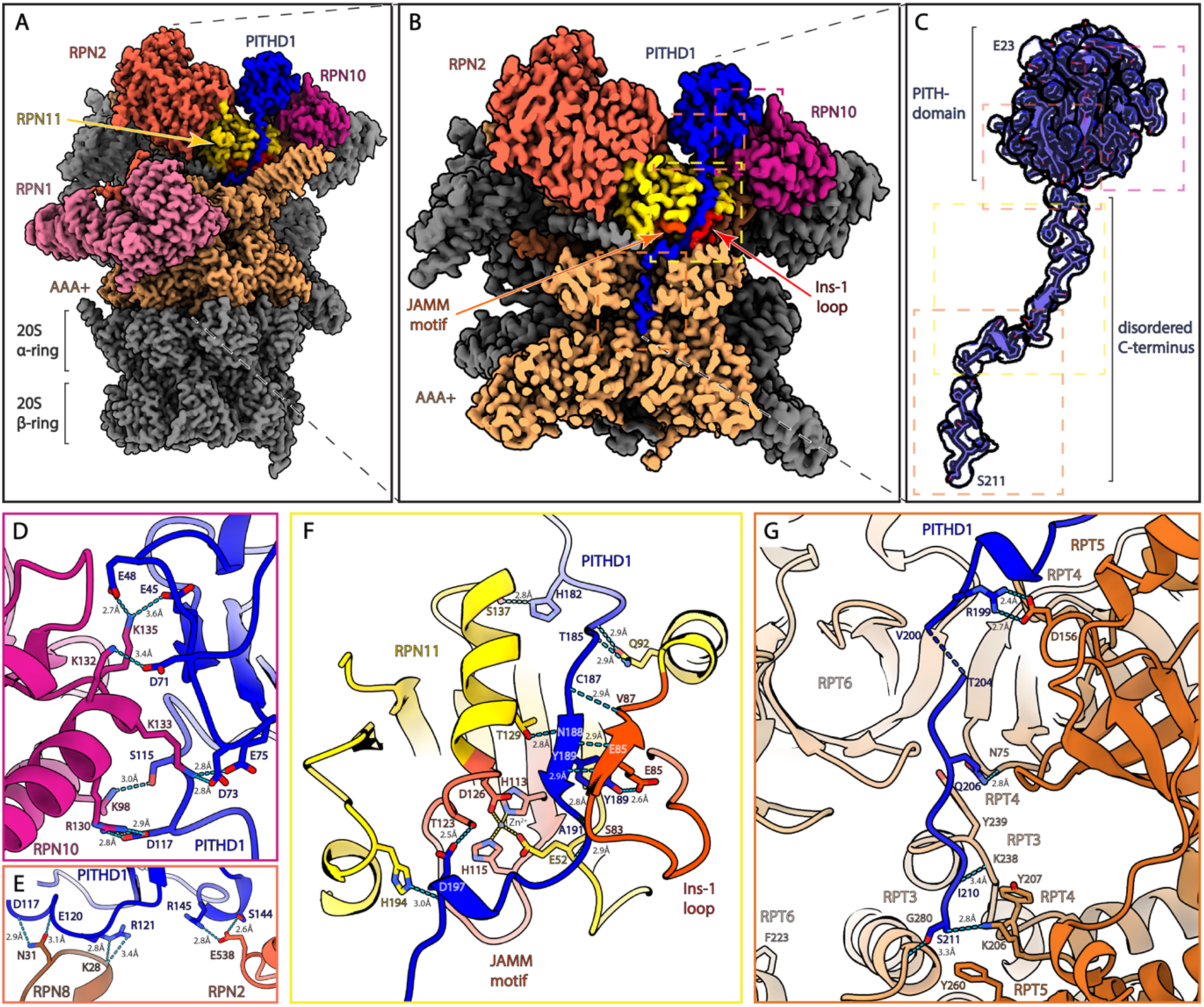
Structure of the PITHD1-26S proteasome complex. (**A**) Consensus map of the reconstituted PITHD1-26S complex. (**B**) A cross-section of the 19S RP reveals interactions of PITHD1’s C-terminus with the active site of the DUB RPN11 and the substrate pore of the ATPase. (**C**) Segmentation of PITHD1 with fitted atomic model. (**D**) Close up of the PITHD1-RPN10 interaction. (**E**) Close-up of the PITHD1-RPN2 and RPN8 interactions. (**F**) Close-up on the PITHD1-RPN11 interaction. (**G**) Close-up on the PITHD1-ATPase interactions.

Our structure reveals that PITHD1 engages multiple functional sites within the 19S regulatory particle through an extensive network of interactions. The globular PITH domain forms specific contacts with three key regulatory components: First, it establishes numerous hydrogen bonds and electrostatic interactions with the main ubiquitin receptor RPN10 (E45/E48-K135, D71-K132, D73/E75-K133, S115-K98, D117-R130; Fig. 2D). Second, it connects to RPN2 through S144/R145-E538 interactions (Fig. 2E). Third, it contacts RPN8 via residue-backbone interactions (D117/E120-N31, R121-K28; Fig. 2E).

While PITHD1’s N-terminus remains largely unresolved, suggesting flexibility, its C-terminus makes interactions that suggest functional importance. Most notably, the C-terminus engages the active site of RPN11, the central proteasomal DUB, through an extensive network of contacts (H182-S137, T185-Q92, C187-V87, N188-T129, Y189-E85, A191-E52/S83, D197-T123/H194; Fig. 2F). This interaction directly blocks the ubiquitin-binding site of RPN11, suggesting that PITHD1 inhibits deubiquitination by restricting substrate access (fig. S5A).

Beyond RPN11, PITHD1’s C-terminus penetrates the central substrate channel, where it forms specific interactions with multiple ATPase subunits. It contacts RPT5 (R199-D156), RPT4 (Q206-N75, S211-K206), and RPT3 (I210-K238, S211-G280; Fig. 2G). Notably, PITHD1 residues I210 and S211 form hydrogen bonds with the pore-1 loops of RPT3 K238 and RPT4 K206, positioning them near three conserved aromatic residues (RPT3 Y239, RPT4 Y207, RPT5 Y260) critical for substrate translocation (*19*, *20*). Despite these extensive interactions, neither wild-type PITHD1 nor a variant with an extended C-terminus (PITHD1+10) is degraded by the proteasome *in vitro* (fig. S5B).

The positioning of PITHD1’s C-terminus in the substrate channel is particularly significant, as this region usually engages unfolded protein domains to initiate translocation toward the 20S core particle (*18*, *20*, *21*). By occupying the substrate channel, PITHD1 would prevent substrate engagement and subsequent degradation (fig. S5A). Collectively, our structural analysis reveals that PITHD1 employs a three-pronged inhibitory mechanism: blocking ubiquitin recognition, preventing deubiquitination, and obstructing substrate translocation.

## PITHD1 acts as an inhibitor of proteasome function

To determine whether PITHD1 directly inhibits proteasome function, we performed degradation assays using purified components. As a model substrate, we used sfGFP(cp8) tagged with an N-terminal K48-linked polyubiquitin chain and a C-terminal unstructured region (fig. S6) (*22*, *23*). The addition of increasing PITHD1 concentrations progressively inhibited substrate processing by the 26S proteasome (Fig. 3A). To validate these findings in cells, we engineered an RKO cell line with doxycycline-inducible PITHD1 expression. Following addition of both an E1 inhibitor (TAK-243) and a DUB inhibitor (PR-619, which spares RPN11), cells with elevated PITHD1 levels showed a markedly reduced clearance rate of K48-ubiquitinated substrates (Fig. 3B).

**Fig. 3.**
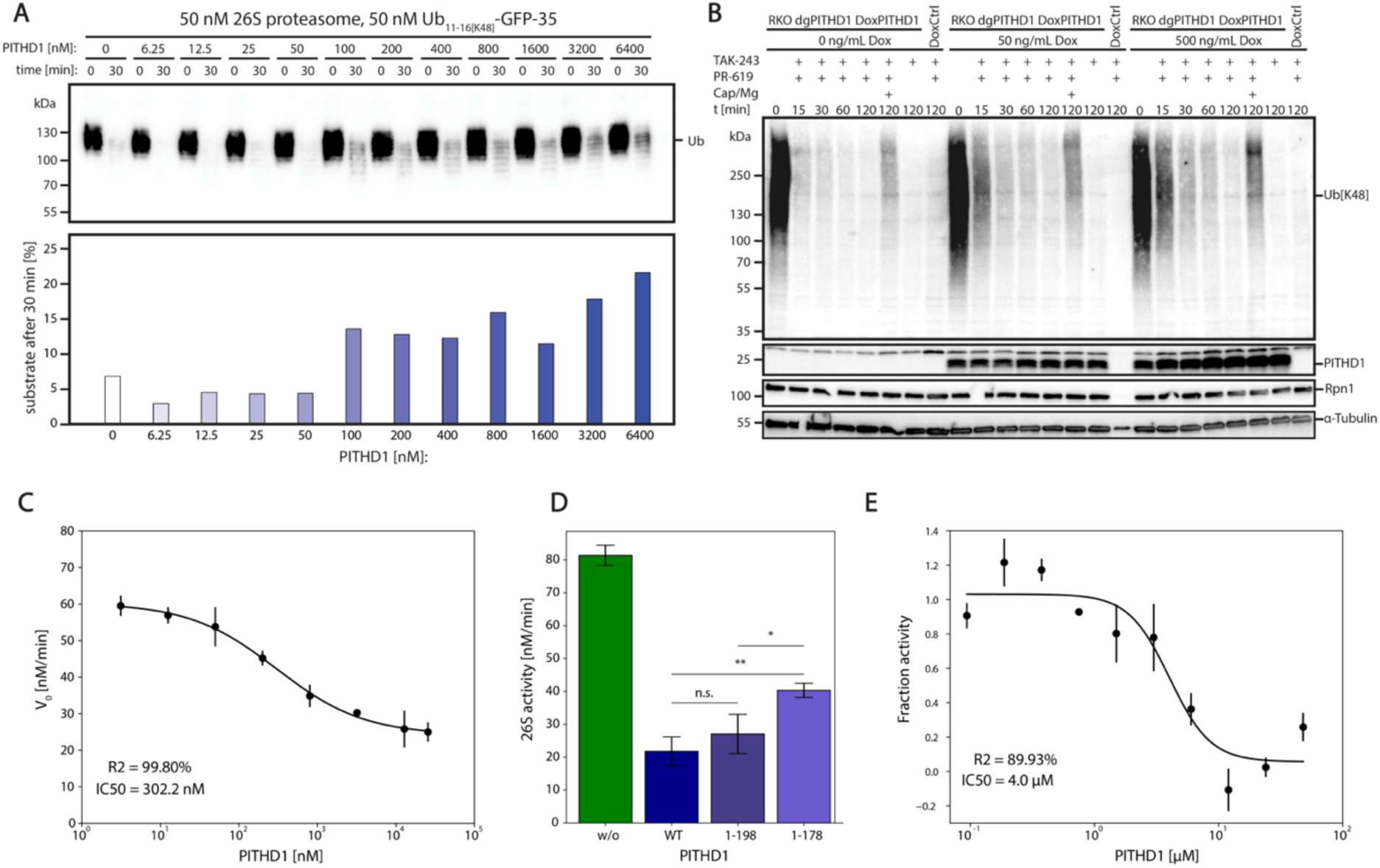
Inhibition of proteasomal activities by PITHD1. (**A**) Degradation assay of poly-Ub[K48]-GFP-35 model substrate, analysed via Western Blot using antibodies against Ub[K48]. Increasing PITHD1 concentration slows down degradation. (**B**) Overexpression of PITHD1 in RKO cells leads to slower degradation of polyubiquitinated species. (**C**) PITHD1-dependent inhibition of K48-ubiquitinated substrate degradation by the 26S proteasome was assessed by measuring GFP fluorescence decay over time (n = 3). (**D**) PITHD1 1-198 and 1-178 mutants show lower inhibitory effect on 26S proteasome activity compared to PITHD1 WT (n = 3, t-test, n.s.: p > 0.05, *: p ≤ 0.05, **: p ≤ 0.01). (**E**) PITHD1 inhibits the DUB activity of the RPN11/RPN8 heterodimer in isolation (n = 3).

Quantitative analysis of the decrease in substrate fluorescence over time allowed us to determine PITHD1’s inhibitory potency. We measured an IC50 of 302 ± 25 nM (Fig. 3C, fig. S7A, B), which falls within the physiological range of cellular PITHD1 concentrations (∼450 nM) (*24*). This suggests that cells can modulate proteasome activity by adjusting PITHD1 levels within their normal dynamic range.

To test structure-based predictions about PITHD1’s inhibitory mechanism, we generated truncation variants. PITHD1 1-198, which cannot enter the central pore but can still interact with RPN11, showed only modest reduction in inhibitory activity. However, PITHD1 1-178, which cannot engage RPN11, displayed substantially decreased inhibition (Fig. 3D, fig S7C, D), while still be able to suppress the proteasome, likely due to its persistent interactions with RPN2, RPN8 and RPN10 via the PITH domain (Fig. 2D, E). These results indicate that PITHD1’s interaction with RPN11 is crucial for its full inhibitory function.

We further dissected PITHD1’s mechanism by examining its effect on isolated RPN8/RPN11 heterodimers (*25*). PITHD1 inhibited RPN11’s deubiquitinating activity with an IC_50_ of approximately 4.0 µM (Fig. 3E, fig. S7E). The higher IC50 compared to full proteasome inhibition suggests that PITHD1’s multiple binding sites act cooperatively to achieve potent inhibition of the intact proteasome.

Together, these biochemical studies demonstrate that PITHD1 functions as a direct proteasome inhibitor through a complex mechanism involving RPN11 inhibition, substrate channel blockade and likely its interaction with RPN2, RPN8 and RPN10. The physiologically relevant inhibitory concentration and the ability to regulate proteasome function in cells support PITHD1’s role as an endogenous proteasome regulator.

## Comparison of PITHD1 to physiological 26S proteasome substrate

To understand how PITHD1 prevents substrate processing, we compared PITHD1-bound and substrate-bound proteasome structures. Previous structural studies have primarily used K63-linked ubiquitin chains (*18*, *20*). However, we reconstituted the 26S proteasome with the physiologically relevant K48-linked ubiquitin chains. Using time-resolved cryoEM (3 s) at elevated pre-freezing temperature (20°C), we captured the transient substrate-bound state preceding deubiquitination at 3.0 Å resolution (fig. S8). This structure revealed that K48-linked ubiquitin occupies the same region on RPN11 as PITHD1 (Fig. 4 A, B).

**Fig. 4.**
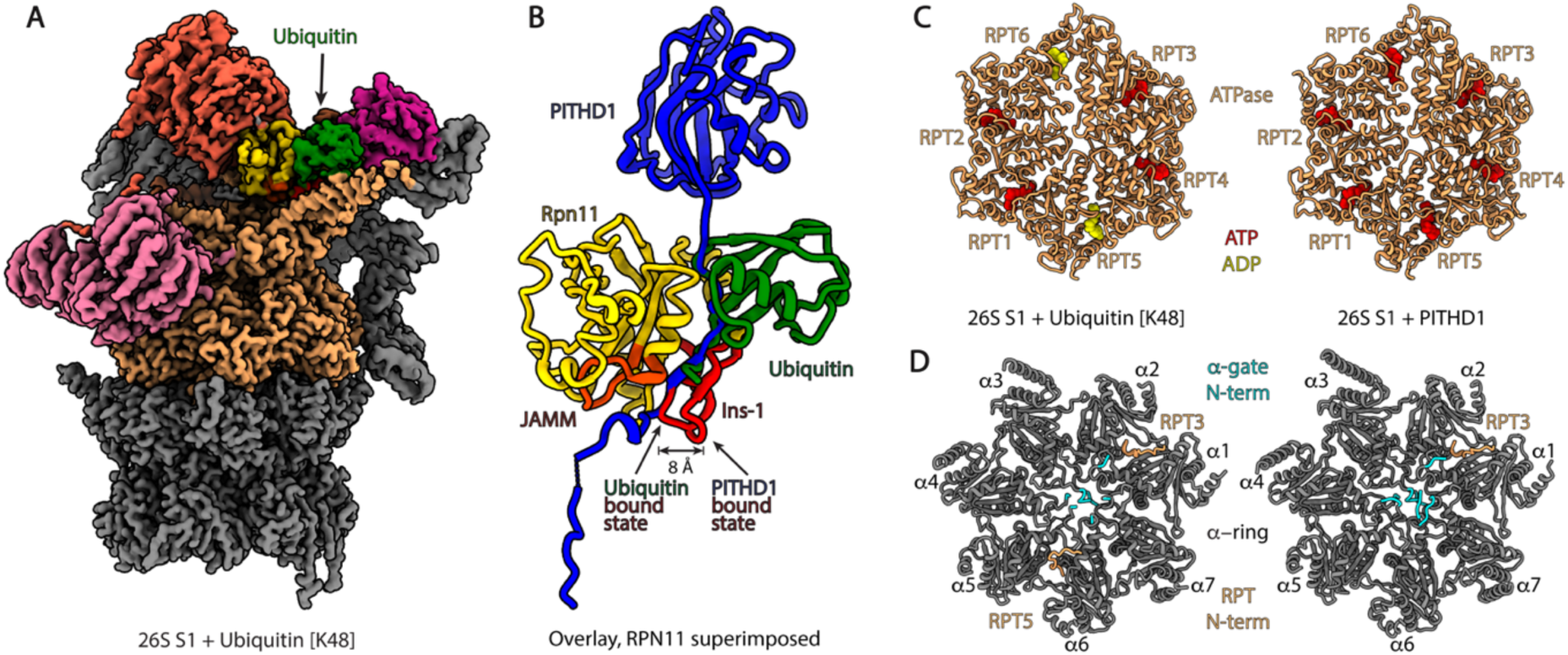
Structural differences between substrate and PITHD1 bound proteasomes. (**A**) Map of K48 ubiquitinated substrate bound proteasome (**B**) Overlay of RPN11 bound to ubiquitin or PITHD1. An outwards displacement of the RPN11 Ins-1 loop can be observed upon the interaction of PITHD1. (**C**) Nucleotide states of the ubiquitin bound and PITHD1 bound proteasome. (**D**) HbYX insertion into α-Ring pockets and α-Ring gate states of the ubiquitin or PITHD1 bound proteasome.

Comparative analysis of PITHD1-bound and substrate-bound states revealed three key conformational differences. First, while substrate-bound proteasomes showed mixed ATP/ADP occupancy, PITHD1-bound proteasomes maintained full ATP occupancy in all six nucleotide-binding pockets (Fig. 4C). This ATP-locked state, combined with PITHD1’s resistance to degradation (fig. S5B), suggests that PITHD1 stabilises a catalytically inactive conformation of the 26S proteasome.

Secondly, PITHD1-bound proteasomes exhibited reduced HbYX motif insertion into the 20S α-ring, accompanied by tighter closure of the 20S α-Ring gate and a decreased chymotrypsin-like activity (Fig. 4D, fig. S9).

Finally, PITHD1 binding repositions RPN11’s Ins-1 loop 8 Å outward relative to the ubiquitin-bound S1 state (Fig. 4B). This outward displacement prevents the formation of the stabilizing β-sheet interaction between the Ins-1 loop and ubiquitin, which is essential for efficient deubiquitination (fig S5A) (*25*). While the β-hairpin formation of the Ins-1 loop upon ubiquitin binding favours a transition to the active S3/S4 states (Movie S1), the outward displacement caused by PITHD1 binding would inhibit this transition by sterically clashing with the RPT4/5 coiled-coil domain (Movie S2).

These structural changes collectively explain how PITHD1 locks the proteasome in an inactive conformation through three coordinated mechanisms: First, it maintains an ATP-bound state that prevents productive substrate engagement. Second, it promotes gate closure in the 20S core particle. Third, it prevents RPN11 activation by displacing the Ins-1 loop. This multi-layered inhibitory strategy ensures a robust suppression of 26S proteasome function during cellular dormancy.

## Binding of PITHD1 to activated proteasomes

To further understand PITHD1’s inhibitory mechanism, we examined its ability to bind activated proteasome states. Treatment with ATPγS typically stabilizes proteasomes in activated S2, S3, and S4 conformations (*26*, *27*). Addition of PITHD1 to ATPγS-treated proteasomes caused a marked redistribution of states (fig. S11A, B). Conformational analysis by 3D classification revealed that PITHD1 increased the S1-state populations by ∼20% while decreasing S2 and S3/4 states by up to 24% and 47%, respectively (fig. S11A, B), demonstrating its ability to drive proteasomes toward inactive conformations.

Unexpectedly, we captured PITHD1 bound 26S proteasomes in both S2-like and an S4 states (Fig. 5 A, B, fig S10). In the S2-like state (2.9 Å resolution), characterized by an RPN1 rotation, PITHD1’s C-terminus remained engaged with RPN11, maintaining the Ins-1 loop in an outward conformation displaced by 11 Å compared to the substrate-engaged S2 state (Fig. 5 A, C, fig. S11C). This state exhibited full ATP occupancy, reduced HbYX motif interactions, and a closed 20S α-ring gate, consistent with an inhibited S1-S2 intermediate (Fig. 5E, fig. S11E).

**Fig. 5.**
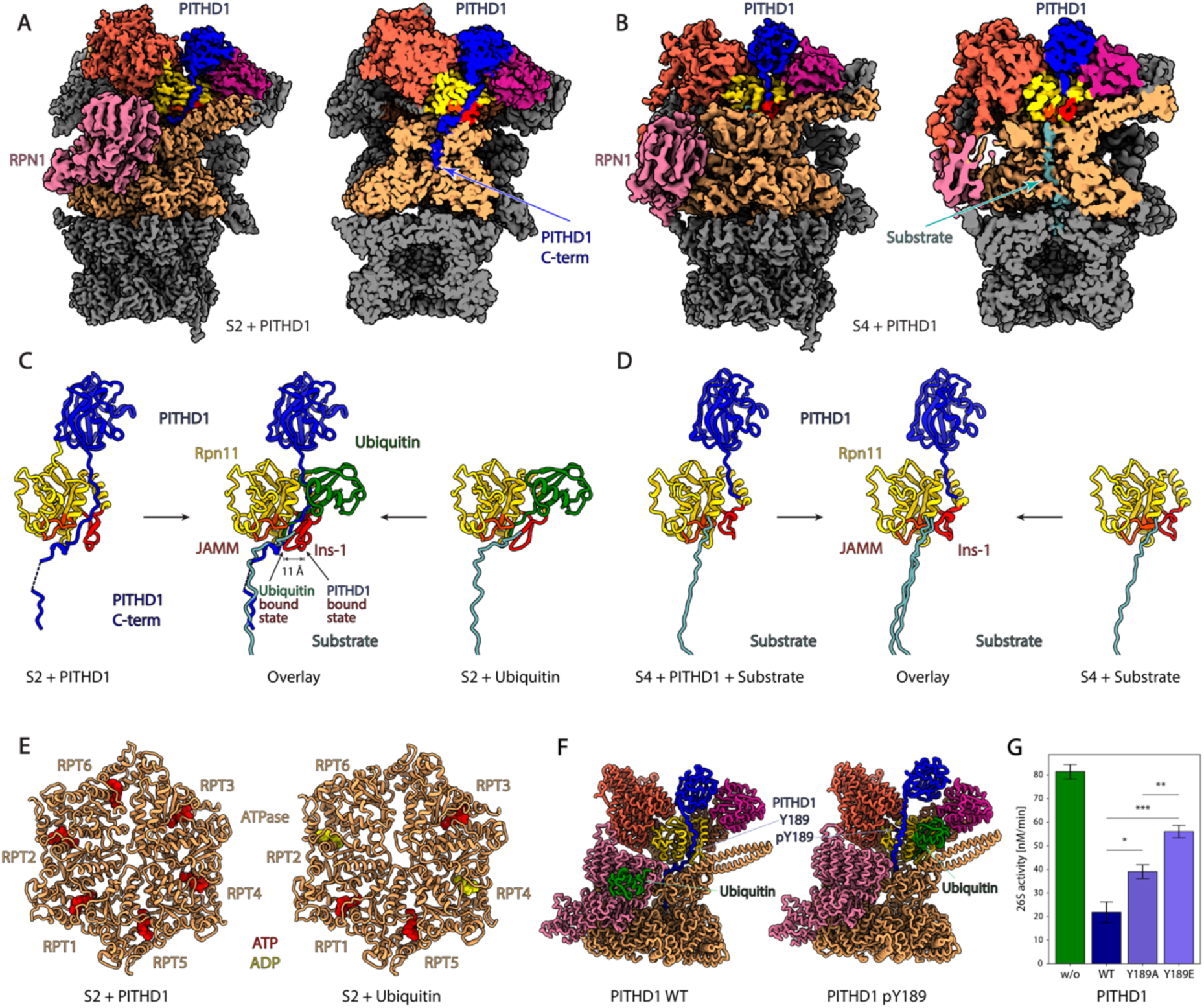
PITHD1 interaction with the activated proteasome. (**A**) Map of PITHD1 bound to the S2 like state. (**B**) Map of PITHD1 bound to the active S4 state. (**C**) Overlay of substrate and PITHD1 bound S2 states. (**D**) Overlay of substrate and PITHD1 bound S4 states. (**E**) Nucleotide occupancy of PITHD1 and Substrate bound S2 states. (**F**) AF3 predictions of functional 19S subunits, ubiquitin and PITHD1 WT or PITHD1 pY189. PITHD1 Y189 phosphorylation facilitates ubiquitin binding. (**G**) Proteasome activity inhibition dependent on PITHD1 WT, Y189A and Y189E mutants (n = 3, t-test, *: p ≤ 0.05, **: p ≤ 0.01, ***: p ≤ 0.001).

In the S4 state (3.5 Å resolution), where RPN11 aligns with the central pore and the ATPase ring engages the 20S CP channel, PITHD1’s C-terminus was disengaged (Fig. 5 B, D). This state showed hallmarks of active proteasomes: a characteristic Ins-1 loop conformation, four HbYX motifs interacting with the α-ring pockets, an open α-ring gate, and a typical nucleotide occupancy pattern (Fig. 5D, fig. S11D, F, I). The presence of an unfolded substrate in the substrate channel in this PITH domain-bound S4 state suggests that, while PITHD1 and ubiquitin binding are mutually exclusive, the interaction of the globular PITH domain alone permits substrate translocation to proceed.

These structures reveal PITHD1’s mechanism of action. It promotes inactive conformations through direct competition with ubiquitin binding, but its release enables the proteasome to resume normal substrate processing. This provides a molecular foundation for understanding PITHD1’s role as a reversible endogenous proteasome inhibitor.

## Regulation of PITHD1 binding through phosphorylation

Detailed analysis of the PITHD1-RPN11 interface revealed that PITHD1’s Y189 residue is deeply embedded in a pocket behind the Ins-1 loop, where it forms specific contacts with RPN11 E85 (Fig. 2F). Database analysis identified Y189 as PITHD1’s most frequently reported phosphorylation site (*28*), suggesting potential regulatory significance. Using AlphaFold3 predictions, we found that phosphorylation of Y189 would create steric and electrostatic conflicts with RPN11’s active site, which could favour the binding of ubiquitin instead (Fig. 5F, Movie S3).

To test this predicted regulatory mechanism, we generated two Y189 variants. A phosphomimetic Y189E mutation and an alanine substitution (Y189A) that eliminates the interaction with RPN11 E85. Both variants showed reduced inhibitory activity in proteasome degradation assays, with the phosphomimetic Y189E displaying a more pronounced effect (Fig. 5G, fig. S11G, H). The stronger impact of Y189E suggests that negative charge at this position actively disrupts PITHD1-proteasome interactions, consistent with our structural predictions.

This phosphorylation-based regulatory mechanism could provide cells with a rapid means to relieve proteasome inhibition. Given that Y189 phosphorylation would specifically disrupt PITHD1’s interaction with RPN11 while potentially leaving other contact points intact, this modification might allow for fine-tuned regulation of PITHD1’s inhibitory function.

## Discussion

The regulated arrest of protein degradation is essential for cellular dormancy, yet its molecular mechanisms have remained unclear. Our discovery of PITHD1 as the first endogenous inhibitor of the 26S proteasome reveals how cells can rapidly and reversibly suppress protein degradation. Through a remarkable triple-lock mechanism, PITHD1 simultaneously blocks ubiquitin recognition, deubiquitination, and substrate translocation – providing more comprehensive proteasome inhibition than other known regulators such as ECM29 or PSMF1. Notably, recent structural studies show that K11-linked ubiquitin chains occupy the same binding site as PITHD1’s globular domain (EMDB-37269, EMDB-37317, EMDB-37334), suggesting that PITHD1 may have evolved to mimic and compete with this specific ubiquitin chain topology, which is essential for cell cycle control.

PITHD1’s widespread presence in cells (*16*) and its regulation by phosphorylation suggest it may serve as a general proteostasis regulator beyond dormancy. While proteasome levels can be adjusted through slow transcriptional responses, PITHD1 phosphorylation may provide a rapid switch for modulating proteasome activity. Further studies examining the kinases and phosphatases that control Y189 phosphorylation state could reveal how cells modulate proteasome activity. This mechanism could be particularly important during cellular state transitions, where quick changes in protein degradation are required.

Outside its function in oocytes, diverse biological roles of PITHD1 are only beginning to emerge. Its requirement for male fertility in both flies (*29*) and mice (*30*) suggests broader functions in cellular differentiation and tissue maintenance.

Our findings establish proteasome inhibition as a key regulatory mechanism in cellular dormancy, complementing known controls on protein synthesis. The discovery that both major cellular machines for protein homeostasis – ribosomes and proteasomes – are subject to specific inhibition during dormancy reveals the sophistication of cellular proteostasis regulation. Understanding how cells coordinate these parallel pathways, and how PITHD1’s activity is regulated in different contexts, represents an important direction for future research. Beyond dormancy, PITHD1’s mechanism, particularly its mimicry of K11 chain binding, suggests new approaches for therapeutic modulation of proteasome activity in diseases ranging from cancer to neurodegeneration.

## Methods

### Zebrafish husbandry

For all experiments TLAB fish were used, which were generated by crossing fish of the TL (Tupfel Longfin) line to fish of the AB line. All samples were collected from age-matched TLAB zebrafish of fish aged between 4-8 months and treated equally. Fish were raised according to standard protocols (28°C; 14/10 h light/dark cycle) and all experiments were conducted according to the Austrian and European guidelines for animal research and approved by the ‘Amt der Wiener Landesregierung’, Magistratsabteilung 58-Wasserrecht (zebrafish protocol MA 58-221180-2021-16).

### Zebrafish sample collection

For the collection of mature zebrafish eggs, female zebrafish were anesthetized in the morning using 0.01% (w/v) Tricaine in fish water (prepared from 25x Tricaine stock in dH_2_O, pH 7.0 – 7.5). After anesthesia the female was transferred to a dry Petri dish, dried off and the eggs expelled from the female using mild pressure from anterior to posterior. Eggs were carefully separated from the female using a paintbrush and the female was returned to fish water for recovery. To prevent activation, eggs were collected in ice-cold sorting medium (90% Leibowitz’s medium, 0.5% w/v BSA and 100 μg/ml G418 sulfate, pH 9.0 with 5 M NaOH; (*31*)) prior to lysis. Each replicate consisted of eggs from two females.

For the collection of ovaries, female zebrafish were euthanized in Tricaine solution until gill activity fully stopped and were subsequently decapitated. Eggs were stripped as described above and discarded. Ovaries were dissected out and any eggs still attached to the ovaries manually removed.

Embryos were collected following mating and kept at 28 °C in E3 medium (5 mM NaCl, 0.17 mM KCl, 0.33 mM CaCl_2_, 0.33 mM MgSO_4_, 10^−5^% Methylene Blue). After reaching the 1h post fertilisation stage, embryos were incubated in 1 mg/mL pronase in E3 medium for dechorionation.

Eggs, ovaries and dechorionated embryos were washed using ice-cold 1x PBS and proteasome buffer (25 mM BisTris pH 6.5, 50 mM KCl, 5 mM MgCl_2_, 10% glycerol, 5 mM ATP, 0.5 mM TCEP) and were subsequently mechanically lysed using pre-cooled dounce homogenizers. For this the ovary and egg samples were homogenized in 400 µl of proteasome buffer and the embryo samples in 250 uL of proteasome buffer with 20 strokes of the loosely fitting pestle A, followed by 20 strokes of the tightly fitting pestle B. Lysates were treated with 1 μL Benzonase prior to proteasome purification.

All samples were collected from age-matched TLAB zebrafish of fish aged between 4-8 months and treated equally.

### Zebrafish Gradients

Lysed ovaries, eggs, and embryos were centrifuged at 10,000 x g for 10 minutes at 4°C to remove debris. To crudely purify 26S proteasomes, the supernatants were subjected to sucrose density gradient ultracentrifugation. Samples were spun at 105,000 x g for 16 hours in a 4 mL tube with proteasome buffer (25 mM Bis-Tris, pH 6.5, 50 mM KCl, 5 mM MgCl₂, 5 mM ATP, 0.5 mM TCEP) and a 10–30% sucrose gradient. Fractions of 200 µL were collected and analyzed by SDS-PAGE and Western blot using anti-RPN1 antibody (ab140675, 1:10,000 dilution) and anti-20S-α1/α2/α3/α5/α6/α7 antibody (sc58412, 1:10,000 dilution) to detect the 26S proteasome.

### Chymotrypsin like activity of Zebrafish proteasomes

Gradient fractions of Zebrafish ovaries, eggs, and embryos containing both Rpn1 and 20S α subunits were pooled and normalized based on 20S α subunit concentrations determined by Western blot (sc58412, 1:10,000 dilution). In triplicates, 20 µL of sample and 10 µL of 30 µM Suc-LLVY-AMC in 26S proteasome activity buffer (25 mM Tris, pH 7.5, 4 mM MgCl₂, 4 mM ATP, 5% glycerol, 1 mM DTT, 2 g/L BSA) were mixed in a microplate (Greiner, 384-well, PS, F-Bottom, Black, Non-Binding, Item No.: 781900). The plate was pre-incubated with 26S activity buffer for 60 minutes at 37°C and dried with compressed air. Fluorescence was measured for 2 hours at 37°C, once per min, using a Synergy H1 (BioTek) microplate fluorescence reader (360 nm excitation, 450 nm emission). In Python, linear regression models were applied to the linear phase of the reaction (60 minutes), and the initial degradation rates ± SD were calculated from the slopes.

### Zebrafish gradients mass spectrometry and AlphaFold2 IP screen

Gradient fractions of Zebrafish ovaries, eggs, and embryos containing both RPN1 and 20S α subunits were pooled, concentrated to 200 µL using Cytiva Vivaspin concentrators (6 mL, 10 k MWCO, 28-9322-96), and buffer-exchanged into proteasome buffer (25 mM Bis-Tris, pH 6.5, 50 mM KCl, 5 mM MgCl₂, 5 mM ATP, 0.5 mM TCEP) using Zeba Spin Desalting Columns (0.5 mL, 7K MWCO, 89882). Further concentration was achieved by acetone precipitation (800 µL pre-cooled acetone at-20°C, overnight incubation, then centrifugation at 21,000 x g for 10 min at 4°C). Pellets were dissolved in 30 µL of 20 mM HEPES and 8 M Urea, pH 7.2 buffer, for tandem MS analysis.

To account for differences in yolk protein composition among samples and compare potential 26S proteasomal cofactors, the dataset was normalized to 26S proteasome subunits. Potential cofactors interacting with the 26S proteasome were identified by filtering for proteins copurified with RPN11-HTBH-tagged 26S proteasomes from HEK293 RPN11-HTBH cells (see below) and analysed via tandem MS. Proteins with a fold change greater than or less than 2-fold in oocytes or eggs relative to embryos were screened using AlphaFold2 against all 26S proteasome subunits. Predictions with an average pTM (predicted template modelling) score above 0.5 and an average ipTM (interface predicted template modelling) score above 0.6 were considered hits (*8*).

To explore their mode of action, AlphaFold 2 hits were further screened against identified binding partners and known 19S RP functional subunits (RPT1-6, RPN1, RPN2, RPN8, RPN10, RPN11) using the AlphaFold3 server (alphafoldserver.com) (*11*).

### PITHD1 conservation in Vertebrates

For the conservation scores, we selected vertebrate sequences from the NCBI protein or UniProt database in blast searches using the human ortholog of PITHD1 (Q9GZP4) and applying highly significant E-value thresholds (<10^-50^) (*32–34*). Orthologous sequences were aligned with mafft (-linsi mode, v7.505, (*35*), and visualized with Jalview (*36*). The per residue sequence conservation scores were calculated with AAcon (v. 1.1., KARLIN method, results normalized with values between 0 and 1, (*37*).

### Mammalian Cell Growth conditions

HEK293 RPN11-HTBH cells expressing hexahistidine, TEV cleavage site, biotinylation site, hexahistidine (HTBH)-tagged PSMD14, a gift from L. Huang (*38*) was grown in 600 mL FreeStyle 293 Expression Medium (Gibco) in 2 L Erlenmeyer flasks. The cells were harvested by centrifugation (15 min, 500 x g, 4°C) at a cell density of 3×10^6^ cells/mL. The pellets were weighed, flash-frozen in liquid nitrogen and stored at-70°C.

### PITHD1 knockout and overexpression in RKO cells

RKO iBFP:Cas9 cells a gift from J. Zuber (*10*) containing pLentiv2-TRE3G-Cas9-P2A-BFP were transduced with dual_sgrna_pithd1_exon1_exon3_thy1.1 modified from Dual-sgRNA_hU6-mU6 (*39*) and packaged in HEK293 LentiX cells. Cas9 expression was induced with 200 ng/mL doxycycline, single cell clones and a batch were FACS sorted for Thy1.1 positive cells by the VBCF optics facility and a clone with a PITHD1 knockout (RKO iBFP:Cas9_dgPITHD1) was identified by Western Blot using Anti-PITHD1 antibody (HPA016936, 1:2,000 dilution). RKO iBFP:Cas9_dgPITHD1 was further transduced with pLentiv2-TRE3G-PITHD1-PGK-GFP-P2A-BlastR which will induce PITHD1 expression upon Doxycycline induction or pLentiv2-TRE3G-Ctrl-PGK-GFP-P2A-BlastR which will not express PITHD1 upon Doxycycline induction, both modified from pLentiv2-TRE3G-Cas9-P2A-GFP-PGK-BlastR (*10*) and packaged in HEK293 LentiX cells. Batches were FACS sorted for GFP-positive cells by the VBCF optics facility. Batch sorted PITHD1 knockout/Dox inducible PITHD1 RKO cells (RKO iBFP:Cas9_dgPITHD1_iGFP:PITHD1) and PITHD1 knockout/Dox inducible Ctrl RKO cells (RKO iBFP:Cas9_dgPITHD1_iGFP:Ctrl) were tested for PITHD1 expression upon the addition of 50 ng/mL or 500 ng/mL doxycycline by Western Blot using Anti-PITHD1 antibody (HPA016936, 1:2,000 dilution).

### Recombinant production of PITHD1 and its variants

N-terminally tagged twinStrep-3C-PITHD1 WT was cloned by inserting E. coli codon-optimized human PITHD1 into TS_pGC-01;02 (Vienna Biocenter Molecular Biology Core Facility) using Gibson cloning. For twinStrep-3C-PITHD1+10, a 30 bp sequence encoding the 10 C-terminal amino acids of Ub-CP8-35ΔK-His6_pET26b (SPAEHHHHHH, a gift from A. Matouschek (*22*, *23*) was cloned downstream to PITHD1’s C-terminus. Variants twinStrep-3C-PITHD1 1-198 and 1-178 were generated via Gibson cloning, and twinStrep-3C-PITHD1 Y189A and Y189E were produced using quick exchange site-directed mutagenesis. All twinStrep-3C-PITHD1 variants were transformed into E. coli Arctic Express and grown in 4 L auto-induction media (10 g/L tryptone, 5 g/L yeast extract, 25 mM (NH4)2SO4, 50 mM KH2PO4, 50 mM Na2HPO4, 0.5% glycerol, 0.05% glucose, 0.2% α-lactose, 1 mM MgSO4) with 100 µg/mL Ampicillin (6 hours at 37°C, overnight at 11°C). Cells were harvested by centrifugation (15 minutes, 4,000 x g, 4°C), resuspended in lysis buffer (25 mM Tris-HCl pH 7.5, 300 mM NaCl, 1 mM DTT, 1× Benzonase, 2× protease inhibitor cocktail [Roche cOmplete]), and lysed using a cell disruptor at 1.5 kbar. Lysates were cleared by centrifugation (1 hour, 18,000 x g, 4°C).

On an ӒKTA pure system (Cytiva), cleared lysates were loaded onto a 5 mL Strep Trap HP column (Cytiva) equilibrated with buffer A_Str_ (25 mM Tris-HCl pH 7.5, 300 mM NaCl, 1 mM DTT). The column was washed with 10 column volumes of buffer AStr, and PITHD1 variants were eluted with buffer B_Str_ (25 mM Tris-HCl pH 7.5, 300 mM NaCl, 1 mM DTT, 5 mM Desthiobiotin). The twinStrep-3C tags were cleaved overnight using 3C protease and further purified via size exclusion chromatography on a HiLoad 16/600 Superdex 75 pg column (Cytiva) using Buffer GF (25 mM Tris-HCl pH 7.5, 100 mM NaCl, 1 mM DTT). Purified PITHD1 and variants were analyzed by Western blot using Anti-PITHD1 antibody (HPA016936, 1:2,000 dilution). Protein concentrations were determined by measuring absorption at 280 nm with respective extinction coefficients.

### 26S proteasome purification

HEK293 cells expressing hexahistidine, TEV cleavage site, biotinylation site, hexahistidine (HTBH)-tagged PSMD14, a gift from L. Huang (*38*) were dounce-homogenized in lysis buffer (25 mM BisTris pH 6.5, 50 mM KCl, 5 mM MgCl2, 10% glycerol, 5 mM ATP, 0.5 mM TCEP, 0.1 mM PMSF, 1x Benzonase, 1% NP40) containing 1× protease inhibitor cocktail (Roche cOmplete). The lysate was cleared by centrifugation (1 hour, 50,000 x g, 4°C) and filtered using Miracloth and 0.45 µm filters.

Using an ӒKTA pure system (Cytiva), the cleared lysate was loaded onto a 1 mL HP Streptavidin column (Cytiva) equilibrated with 26S proteasome buffer (25 mM BisTris, pH 6.5, 50 mM KCl, 5 mM MgCl₂, 10% glycerol, 5 mM ATP, 0.5 mM TCEP). The column was washed with 10 column volumes (CV) of 26S proteasome buffer. Proteasomes were cleaved from the column overnight using GST-tagged TEV protease.

To remove GST-tagged TEV protease, a 5 mL GSTrap HP column (Cytiva) equilibrated with 26S proteasome buffer was used. Cleaved 26S proteasomes were eluted with 26S proteasome buffer, and GST-TEV protease was eluted separately with GST-elution buffer (25 mM BisTris, pH 6.5, 50 mM KCl, 5 mM MgCl₂, 10 mM GSH, adjusted to pH 7.0).

Western blotting was performed to detect RPN1 and GST in the 26S proteasome and GST elutions using anti-RPN1 antibody (ab140675, 1:10,000 dilution) and anti-GST antibody (RPN1236, 1:10,000 dilution). 26S proteasome concentrations were quantified using the Bradford assay with a BSA standard curve. Their degradation activity was tested with poly-Ub[K48]-GFP-35 (see below), and their composition was analyzed by tandem MS.

### Purification of UBA1

E. coli BL21(DE3) cells were transformed with the plasmid Ube1/PET21d (AddGene Plasmid #34965) and grown in 6 L auto-induction media with 100 µg/mL Ampicillin (6 hours at 37°C, overnight at 18°C). Cells were harvested by centrifugation (15 minutes, 4,000 x g, 4°C), resuspended in lysis buffer (20 mM Tris-HCl, pH 7.5, 300 mM NaCl, 0.5 mM TCEP, 1× Benzonase, 3× protease inhibitor cocktail [Roche cOmplete]), and lysed using a cell disruptor at 1.5 kbar. Lysates were cleared by centrifugation (1 hour, 18,000 x g, 4°C).

Using an ӒKTA pure system (Cytiva), cleared lysates were loaded onto two 5 mL FF Crude HisTrap columns (Cytiva) equilibrated with buffer AHis (20 mM Tris-HCl, pH 7.5, 300 mM NaCl, 0.5 mM TCEP). The columns were washed with 10 column volumes (CV) of buffer A_His_ containing 5% buffer B_His_ (20 mM Tris-HCl, pH 7.5, 300 mM NaCl, 0.5 mM TCEP, 500 mM Imidazole), and His-UBA1 was eluted with buffer A_His_ containing 50% B_His_.

Eluted UBA1 was diluted 10× into buffer A_ResQ_ (50 mM Tris-HCl, pH 7.5, 50 mM NaCl, 0.5 mM TCEP), loaded onto a 6 mL ResQ column (Cytiva) equilibrated with buffer A_ResQ_, and washed with 5 CV of buffer A_ResQ_. His-UBA1 was eluted using a gradient of buffer A_ResQ_ and 0–37% buffer B_ResQ_ (50 mM Tris-HCl, pH 7.5, 2 M NaCl, 0.5 mM TCEP).

His-UBA1 was further purified by size exclusion chromatography on a HiLoad 16/600 Superdex 200 pg column (Cytiva) using Buffer GF (50 mM HEPES-KOH, pH 7.4, 100 mM NaCl, 0.5 mM TCEP, 10% glycerol). Purified His-UBA1 was analyzed by SDS-PAGE, tested for activity in ubiquitination assays, and quantified by measuring absorption at 280 nm using the respective extinction coefficient.

### Purification of SuperE2 Enzyme

E. coli Rosetta2 cells were transformed with a plasmid encoding the E3 RING–E2 fusion protein SuperE2 (a gift from N. G. Brown) and grown in 6 L auto-induction media with 100 µg/mL Ampicillin and 17 µg/mL Chloramphenicol (6 hours at 37°C, overnight at 18°C). Cells were harvested by centrifugation (15 minutes, 4,000 x g, 4°C), resuspended in lysis buffer (25 mM Tris-HCl, pH 8.0, 300 mM NaCl, 0.5 mM TCEP, 1× Benzonase, 2× protease inhibitor cocktail [Roche cOmplete]), and lysed using a cell disruptor at 1.5 kbar. Lysates were cleared by centrifugation (1 hour, 18,000 x g, 4°C).

Using an ӒKTA pure system (Cytiva), cleared lysates were loaded onto a 5 mL FF Crude HisTrap column (Cytiva) equilibrated with buffer AHis (25 mM Tris-HCl, pH 8.0, 300 mM NaCl, 0.5 mM TCEP). The column was washed with 10 column volumes (CV) of buffer A_His_ containing 5% buffer B_His_ (25 mM Tris-HCl, pH 8.0, 300 mM NaCl, 0.5 mM TCEP, 500 mM Imidazole). His-SuperE2 was eluted using buffer AHis with 50% BHis.

His-SuperE2 was further purified by size exclusion chromatography using a HiLoad 16/600 Superdex 75 pg column (Cytiva) and Buffer GF (25 mM Tris-HCl, pH 8.0, 100 mM NaCl, 0.5 mM TCEP). Purified His-SuperE2 was analyzed by SDS-PAGE, and its activity was tested in ubiquitination assays. Protein concentrations were determined by measuring absorption at 280 nm using the respective extinction coefficient. The protein’s activity and specificity for K48-linked ubiquitin chain formation were tested using ubiquitin WT and the K48R mutant (see below).

### Purification of wild-type Ubiquitin

E. coli BL21(DE3) cells were transformed with plasmid pET3a-UbWT (AddGene Plasmid #26685) and grown in 4 L auto-induction media with 50 µg/mL Kanamycin (6 hours at 37°C, overnight at 18°C). Cells were harvested by centrifugation (15 minutes, 4,000 x g, 4°C), and pellets were resuspended in Ubiquitin Buffer (50 mM Tris, pH 7.5, 300 mM NaCl) containing one cOmplete protease inhibitor cocktail tablet and Benzonase. Cells were lysed using a French Press at 1.5 kBar and centrifuged for 1 hour at 40,000 x g.

The supernatant was adjusted to pH 4.5 with concentrated acetic acid and centrifuged (30 minutes, 15,000 x g, 4°C). The resulting supernatant was dialyzed overnight into 25 mM sodium acetate (NaAc), pH 4.5, concentrated, and loaded onto a Resource S column (Cytiva) equilibrated with 25 mM NaAc, pH 4.5. The column was washed with 25 mM NaAc, pH 4.5, 92.5 mM NaCl, and ubiquitin was eluted with 25 mM NaAc, pH 4.5, 250 mM NaCl.

Peak fractions (A280nm) were concentrated using a 5 kDa cut-off concentrator (Cytiva) and further purified on a Superdex 75 16/60 column (Cytiva) equilibrated with buffer (25 mM HEPES, pH 8, 50 mM NaCl). Fractions with a peak at ∼8.6 kDa were analyzed by SDS-PAGE, and those containing ubiquitin were pooled. Protein concentrations were measured using absorption at 280 nm and the respective extinction coefficient.

### Purification of Ubiquitin K48R

E. coli BL21(DE3) cells were transformed with plasmid UBK48R_pGEX (a gift from N. G. Brown), encoding GST-TEV-Ubiquitin K48R, and grown in 6 L auto-induction media supplemented with 100 µg/mL Ampicillin (6 hours at 37°C, overnight at 18°C). Cells were harvested by centrifugation (15 minutes, 4,000 x g, 4°C), and the pellet was resuspended in lysis buffer (25 mM Tris-HCl, pH 8.0, 300 mM NaCl, 1 mM DTT, 1× Benzonase, 1× protease inhibitor cocktail [Roche cOmplete]). The cells were lysed using a cell disruptor at 1.5 kbar, and lysates were cleared by centrifugation (1 hour, 18,000 x g, 4°C).

The cleared lysate was loaded onto a 5 mL GSTrap HP column (Cytiva) equilibrated with buffer A_GST_ (25 mM Tris-HCl, pH 8.0, 300 mM NaCl, 1 mM DTT) using an ӒKTA pure system (Cytiva) and washed with 20 column volumes (CV) of buffer A_GST_. GST-TEV-Ub K48R was eluted with buffer B_GST_ (25 mM Tris-HCl, pH 8.0, 300 mM NaCl, 1 mM DTT, 10 mM GSH).

The GST-TEV tag was cleaved from ubiquitin K48R overnight at 4°C using TEV protease. Cleaved ubiquitin K48R was diluted 10× into buffer A_ResQ_ (25 mM Tris-HCl, pH 8.0, 30 mM NaCl, 1 mM DTT), loaded onto a 6 mL ResQ column (Cytiva) equilibrated with buffer A_ResQ_, and washed with 5 CV of buffer A_ResQ_. Ubiquitin K48R was eluted with a gradient from buffer A_ResQ_ and 0– 37% B_ResQ_ (25 mM Tris-HCl, pH 8.0, 30 mM NaCl, 1 mM DTT, 2 M NaCl).

Protein concentration was determined by absorption at 280 nm using the respective extinction coefficient.

### Purification of Ub-GFP-35

E. coli BL21(DE3) cells were transformed with a plasmid encoding N-terminal Ubiquitin, central sfGFP(cp8), and a C-terminal unfolded domain comprising the first 35 amino acids of cytochrome b2 followed by a hexahistidine tag (Ub-GFP-35), a gift from A. Matouscheck (*22*, *23*). The cells were grown in 4 L auto-induction media supplemented with 50 µg/mL Kanamycin (6 hours at 37°C, overnight at 18°C). Cells were harvested by centrifugation (15 minutes, 4,000 x g, 4°C), and pellets were resuspended in lysis buffer (25 mM Tris-HCl, pH 8.0, 300 mM NaCl, 0.5 mM TCEP, 1× Benzonase, 2× protease inhibitor cocktail [Roche cOmplete]). The cells were lysed using a cell disruptor at 1.5 kbar, and lysates were cleared by centrifugation (1 hour, 18,000 x g, 4°C).

The cleared lysates were loaded onto a 5 mL FF Crude HisTrap column (Cytiva) equilibrated with buffer A_His_ (25 mM Tris-HCl, pH 8.0, 300 mM NaCl, 0.5 mM TCEP) using an ӒKTA pure system (Cytiva). The column was washed with 10 column volumes (CV) of buffer AHis containing 5% buffer B_His_ (25 mM Tris-HCl, pH 8.0, 300 mM NaCl, 0.5 mM TCEP, 500 mM Imidazole), and Ub-GFP-35 was eluted with buffer A_His_ containing 50% B_His_.

Ub-GFP-35 was further purified by size exclusion chromatography using a HiLoad 16/600 Superdex 200 pg column (Cytiva) and Buffer GF (25 mM Tris-HCl, pH 8.0, 100 mM NaCl, 0.5 mM TCEP). The purified protein was analysed by SDS-PAGE, and concentrations were determined by absorption at 280 nm using the respective extinction coefficient.

### Purification of Yeast Rpn8/11

*Sc* Rpn8/Rpn11 and the Ubiquitin substrate were expressed and purified from E. coli and the Ubiquitin substrate was attached to a TAMRA maleimide using cysteine chemistry as previously described (*25*, *40*) All proteins were purified and assayed in GF buffer (50 mM HEPES 7.6, 50 mM NaCl, 50 mM KCl, 10 mM MgCl_2_, 5% glycerol).

### In situ 26S proteasome degradation/deubiquitination assay

Batch sorted RKO iBFP:Cas9_dgPITHD1_iGFP:PITHD1 and RKO iBFP:Cas9_dgPITHD1_iGFP:Ctrl cells were seeded in 6-well plates with 2 mL RPMI medium supplemented with 0, 50, or 500 ng/mL doxycycline and grown to 70% confluency. At 0, 15, 30, 60, and 120 minutes after adding 1 µM E1 inhibitor (TAK-243) and 20 µM panDUB inhibitor (PR619), cells were washed with 2 mL PBS, collected by trypsinization, and centrifuged (5 minutes, 500 x g, 4°C). The pellets were lysed using RIPA buffer (50 mM Tris, pH 8, 150 mM NaCl, 1% Triton, 0.5% sodium deoxycholate, 0.1% SDS, 1 mM PMSF).

Control samples included cells treated with 10 µM Capzimin, 10 µM Mg132, or without PR619, along with RKO iBFP:Cas9_dgPITHD1_iGFP:Ctrl cells at each doxycycline concentration. Samples were analysed by Western blot using Anti-diUbiquitinK48 (AB_1587578, 1:2,000 dilution), Anti-PITHD1 (HPA016936, 1:2,000 dilution), Anti-RPN1 (ab140675, 1:10,000 dilution), Anti-α-Tubulin (T9026, 1:10,000 dilution), and Anti-20S-α1/α2/α3/α5/α6/α7 (sc58412, 1:10,000 dilution) antibodies.

### 26S proteasome PITHD1 reconstitution and cryoEM data collection

#### Sucrose gradient ultracentrifugation in ATP or ATPγS

300 pmol of purified 26S proteasome, with or without 3 µmol PITHD1, was incubated for 60 minutes on ice. The samples were loaded into a 4 mL SW 60 Ti ultracentrifugation tube (Beckman Coulter) containing a 10–30% sucrose gradient in 26S proteasome buffer. For the sample with PITHD1, the gradient buffer contained 4 mM ATP and for the sample without PITHD1, the gradient buffer contained 4 mM ATPγS. The samples were ultracentrifuged in an Optima XE Ultracentrifuge (16 hours, 105,000 x g, 4°C) and fractionated into 200 µL fractions. Fractions were analyzed by Western blot using anti-RPN1 antibody (ab140675, 1:10,000 dilution) and anti-PITHD1 antibody (HPA016936, 1:2,000 dilution).

Peak fractions of 26S proteasomes with PITHD1 in 4 mM ATP were supplemented with 1 µM PITHD1 and incubated on ice for 1 hour. For the peak fraction without PITHD1 in 4 mM ATPγS, 1 µM PITHD1 or PITHD1+10 was added. Control samples without PITHD1 received no additional PITHD1.

#### CryoEM grid preparation

Immediately before grid preparation, all 26S proteasome samples were buffer exchanged into 26S proteasome buffer without glycerol in 4 mM ATP or ATPγS using Zeba Spin Desalting Columns (0.5 mL, 7K MWCO, 89882).

On a Leica EM GP (4°C, 75% humidity), 3 µL of the samples were applied to a holey carbon R3.5/1 Cu200 grid (Quantifoil) covered with a 2 nm continuous carbon foil, plasma-cleaned using an SCD005 sputter coater (Bal-Tec, 60 s, 25 mA). The grids were incubated for 30 seconds, blotted for 2 seconds (Whatman grade 1) using the Leica EM GP automated blotting feature, and plunge-frozen into liquid ethane.

#### Time-resolved cryoEM grid preparation

On an in-house built manual plunge freezing device at RT (20°C, 40% humidity), 1 µL of 10 µM 11-16Ub_[K48]_-GFP-35 was pipetted on a holey carbon R2/2 Cu200 grid (Quantifoil) that was plasma cleaned by a SCD005 sputter coater (Bal-Tec, 60 s, 25 mA). 3 µL of 666 nM 26S proteasome was added, mixed by rapid pipetting, followed by immediate blotting (Whatman grade 1) and plunge freezing into liquid ethane. The total reaction time from mixing to freezing was measured by an assistant with a stopwatch and was 3 s.

#### CryoEM screening and data collections

CryoEM grids were screened on a 200 kV Glacios TEM (Thermo Fisher) at the VBCF EM facility. Grids with good ice quality and intact 26S proteasome particles were selected for data collection on 300 kV Titan Krios TEM instruments.

For the 26S proteasome reconstituted with PITHD1 in ATP and samples without PITHD1, with PITHD1, or with PITHD1+10 in ATPγS, data were collected on a 300 kV Titan Krios G4 TEM (Thermo Fisher) at the IMP. This instrument was equipped with an E-CFEG electron source, a Selectris X imaging filter (Thermo Fisher) with a 10 eV slit width, and a Falcon 4i direct electron detector. Data collection was performed using EPU (Thermo Fisher) with 40 frames per exposure, a pixel size of 0.951 Å²/px, a total dose of 40 e⁻/Å², and a defocus range of −0.6 to −2 µm in −0.2 µm increments.

The time-resolved cryoEM sample was collected on a 300 kV Titan Krios G3i TEM (Thermo Fisher) at ISTA. This instrument was equipped with an X-FEG electron source, a BioQuantum energy filter (Gatan) with a 20 eV slit width, and a K3 direct electron detector (Gatan). Data collection was also performed with EPU, using 40 frames per exposure, a pixel size of 1.07 Å²/px, a total dose of 40 e⁻/Å², and a defocus range of −1.0 to −2 µm in −0.2 µm increments.

#### CryoEM Image processing

All datasets were pre-processed using CryoSPARC v4.6 (*41*), including Patch Motion Correction, Patch CTF Estimation, Curate Exposures, Particle Template Picking, Particle Extraction, 2D Class Cleaning, Ab-initio Reconstruction, and Volume Alignment Tool. Heterogeneity sorting was performed with CryoSPARC v4.6 (Heterogeneous Refinement and 3D Classification)(*41*), RELION v4.0 (3D Classification) (*42*), or cryoDRGN v0.3.4 (Heterogeneous Reconstruction) (*43*). Final 3D refinements were conducted using CryoSPARC v4.6 (Homogeneous Refinement and Local Refinement) (*41*).

For each 26S proteasome state, seven local-refined maps with overlapping regions were reconstructed using ChimeraX v1.8 (*44*). Each locally refined volume (#2-8) was first aligned to the homogeneous consensus map (#1) with the commands “fitmap #2 in #1” to “fitmap #8 in #1” and then combined using the command “vop max #2-8 ongrid #1”. The reconstituted map was post-processed with DeepEMhancer. For further details, see the supplementary material.

#### CryoEM model building

For the 26S proteasomes with PITHD1 in the S1 or S2-like state or with ubiquitin in the S1 state PDB model 6MSD, and for the 26S proteasomes with PITHD1 in the S4 state PDB model 6MSJ was used as an initial model (*20*). The AF2 predicted model for PITHD1 was added using ChimeraX v1.8, ISOLDE v1.8 and Coot v0.9.8.93 (*44–46*).

After an initial refinement with Phenix, several 19S proteasome subunits (SEM1, PSMD2, PSMD3, PSMD6, PSMD8, PSMD13) that did not fit well were replaced by AF2 predicted models using ChimeraX v1.8, ISOLDE v1.8 and Coot v0.9.8.93 (*44–46*) The complete model was refined iteratively using Phenix, with outliers corrected using Coot v0.9.8.93 and ISOLDE v1.8 (*45*, *46*)

#### Preparation of K48 ubiquitinated substrate

Ubiquitination reactions to form K48-linked ubiquitin chains on Ub-GFP-35 were set up overnight (ON) at 30°C. The reactions contained 1.5 µM UBA1, 1 µM SuperE2, 10 mM ATP-MgCl₂, 1 mM Ubiquitin, and 100 µM Ub-GFP-35 in ubiquitination buffer (20 mM HEPES, pH 7.4, 50 mM NaCl). Negative control reactions were performed either without Ub-GFP-35, without ATP, or with Ubiquitin K48R. All reactions were analysed using SDS-PAGE.

To separate the ubiquitinated species according to their chain length, size exclusion chromatography was performed using a HiLoad 16/600 Superdex 200 pg (Cytiva) with Buffer GF (25 mM Tris HCl pH 8.0, 100 mM NaCl, 0.5 mM TCEP). The fractions were analysed by SDS-PAGE and protein concentrations were determined according to their fluorescence with a Ub-GFP-35 standard curve. The degradation of poly-Ub_[K48]_-GFP-35 (≥ 4 ubiquitin moieties) by the 26S proteasome was tested by degradation assays.

#### 26S proteasome degradation assays

To assess the effect of PITHD1 on 26S proteasome degradation of a substrate containing a K48-linked ubiquitin chain, degradation assays were set up with 50 nM 26S proteasome, 50 nM poly-Ub_[K48]_-GFP-35, and varying concentrations of PITHD1 in 26S proteasome activity buffer (25 mM Tris, pH 7.5, 4 mM MgCl₂, 4 mM ATP, 5% glycerol, 1 mM DTT, 2 g/L BSA). Mix1 contained 100 nM poly-Ub[K48]-GFP-35 in 26S proteasome activity buffer, and Mix2 contained 100 nM 26S proteasome with or without varying concentrations of PITHD1. Both mixes were incubated for 60 minutes on ice before they were used in the assay.

#### Western Blot analysis

For Western blot analysis, reactions were initiated at 37°C by mixing 25 µL of Mix2 (100 nM 26S proteasome, 0 - 12.8 µM) with 25 µL of Mix1 (100 nM 11-16Ub[K48]-GFP-35). Aliquots were taken at 0 and 30 minutes, and the reactions were stopped using SDS loading dye. Samples were analyzed by Western blot using an Anti-Ubiquitin antibody (sc-8017, 1:2,000 dilution) and an Anti-PITHD1 antibody (HPA016936, 1:2,000 dilution). Bands were quantified using FIJI, and the residual substrate at 30 minutes was normalized to the 0-minute timepoint.

#### Micoplate fluorescence assays

To measure fluorescence changes over time, the reactions were performed in a microplate fluorescence reader. In triplicates, 25 µL of Mix2 (100 nM 26S proteasome, 0 - 51.2 µM PITHD1), 50 µL blanks (26S activity buffer only), and 50 µL substrate-only samples (50 nM 11-16Ub[K48]-GFP-35 in 26S proteasome activity buffer) were pipetted into a microplate (Greiner, 384 well, PS, F-Bottom, Black, Non-Binding, Item No.: 781900) pre-incubated with 26S activity buffer for 60 minutes at 37°C and dried with compressed air. Using a Synergy H1 (BioTek) microplate fluorescence reader preheated to 37°C, the substrate-only samples were used for automated z-height autofocus and to set the detector gain for 488 nm excitation and 520 nm emission. Reactions were initiated by adding 25 µL of Mix1 (100 nM poly-Ub[K48]-GFP-35) via the integrated dispenser, and fluorescence was measured every minute for 3 hours.

Linear regression models were fitted to the linear phase of the reaction (30 minutes) using Python. Initial degradation rates (V0 [nM/min]) were calculated, and the averages ± SD were plotted on a semi-log graph with a linear V0 scale and a log10-transformed PITHD1 concentration scale. A sigmoidal curve was fitted, and the inflection point of the model was used to calculate the IC50 concentration.

#### PITHD1 dependent Chymotrypsin like activity of the 26S proteasome

To assess the effect of PITHD1 on the 20S α-ring gate closure, chymotrypsin like activity reactions were performed. In triplicates, a mixture of 20 µL of 75 nM 26S proteasome and 10 µL of 30 µM Suc-LLVY-AMC in 26S proteasome activity buffer (25 mM Tris, pH 7.5, 4 mM MgCl₂, 4 mM ATP, 5% glycerol, 1 mM DTT, 2 g/L BSA) was prepared in a microplate (Greiner, 384 well, PS, F-Bottom, Black, Non-Binding, Item No.: 781900). The plate was pre-incubated with 26S activity buffer for 60 minutes at 37°C and dried with compressed air.

Fluorescence was measured every minute for 12 hours using a Synergy H1 (BioTek) microplate fluorescence reader preheated to 37°C, with 360 nm excitation and 450 nm emission. Linear regression models were applied to the linear portion of the reaction (60 minutes) using Python, and initial degradation rates (slopes) ± SD were calculated.

#### Rpn11/8 deubiquitination assays

S. cerevisiae Rpn11 DUB activity was measured using Ubiquitin-5-carboxy tetramethylrhodamine (Ub-TAMRA) as a substrate, as previously described (*25*, *40*). Sc Rpn8/Rpn11 and the Ub-TAMRA substrate were expressed and purified from E. coli, and TAMRA was attached to the ubiquitin substrate via maleimide-cysteine chemistry.

In GF buffer (50 mM HEPES, pH 7.6, 50 mM NaCl, 50 mM KCl, 10 mM MgCl₂, 5% glycerol), 0.5 µM Sc Rpn8/11 was mixed with 50 µM Ub-TAMRA, with or without a two-fold serial dilution of PITHD1 (48 µM to 93.75 nM). Fluorescence polarization (FP) of the TAMRA dye, as it was liberated from Ub-TAMRA, was monitored using a Clariostar Plus (BMG Labtech) with **λ**_ex_ = 540 nm (bandpass 20 nm) and **λ**_em_ = 590 nm (bandpass 20 nm).

The slope of FP loss was calculated using Prism and Python. Degradation rates (V0) were averaged ± SD and plotted on a semi-log graph with a linear V0 scale and a log10-transformed PITHD1 concentration scale. A sigmoidal curve was fitted, and the inflection point of the curve was used to calculate the IC50 concentration.

## Data availability

All data supporting the findings of this study are available within the article, extended data, Supplementary information or the associated data tables. Original raw data will be provided upon request for all experiments, including supporting information. CryoEM raw data was depositied in the EMPIAR database with the accession XXX. The cryoEM density maps have been deposited into the Electron Microscopy Data Bank under accession codes EMD-XXXXX Model coordinates have been deposited into the PDB under accession numbers XXXX.

## Authors Contributions

S.J.A. and J.R. prepared the zebrafish samples for the MS analyses. S.J.A and D.K. generated the cell lines and performed cellular assays. E.R. conducted the MS analyses. S.J.A and A.S performed the homology assessments. S.J.A, I.G. and K.D. purified proteins and conducted the activity assays. S.J.A. prepared cryoEM samples. S.J.A, I.G. and H.K. collected cryoEM data. S.J.A processed the cryoEM data. S.J.A and D.K. did the model building. D.H., A.M. and A.P. supervised this study and devised the methods. S.J.A and D.H wrote the manuscript with input from all authors.

## Acknowledgements

We thank all members of the Haselbach for their great support and scientific discussions, the Electron Microscopy facility of the Vienna BioCenter, in particular Thomas Heuser; We are thankful to the Peters, Plaschka, Clausen and Zuber group for sharing chemicals and lab equipment. This work was supported by the EM Facility and the Protein Technologies Facility at the Vienna BioCenter Core Facilities (VBCF), Proteomics analyses were performed by the Proteomics Facility at IMP/IMBA/GMI using the VBCF instrument pool, in particular Otto Hudecz.

## Supplementary Data

**Figure S1.**
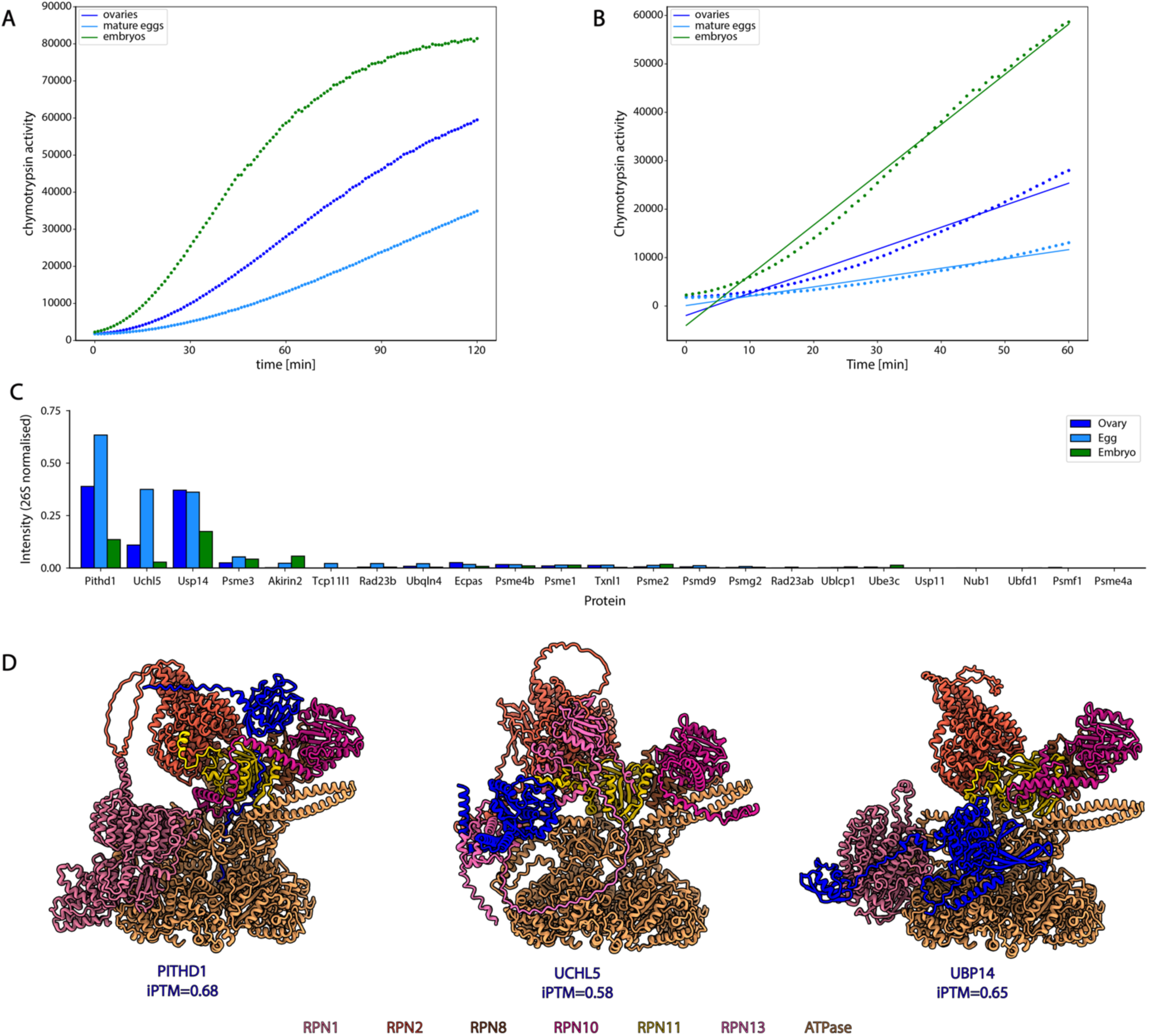
Investigation of zebrafish ovary, egg and embryo 26S proteasomes. (**A**) Chymotrypsin activities of normalised zebrafish 26S proteasome gradient samples, determined using Suc-LLVY-AMC. (**B**) Linear fits of (**A**) to determine the initial degradation rates. (**C**) Mass spectrometry intensities of all potential 26S proteasomal cofactors in the three zebrafish samples identified in the HEK293 26S proteasome AlphaFold interaction screen. (**D**) AF3 predictions of the functional 19S RP subunits and the most abundant potential 26S proteasomal interactors of the AlphaFold interaction screen.

**Figure S2.**
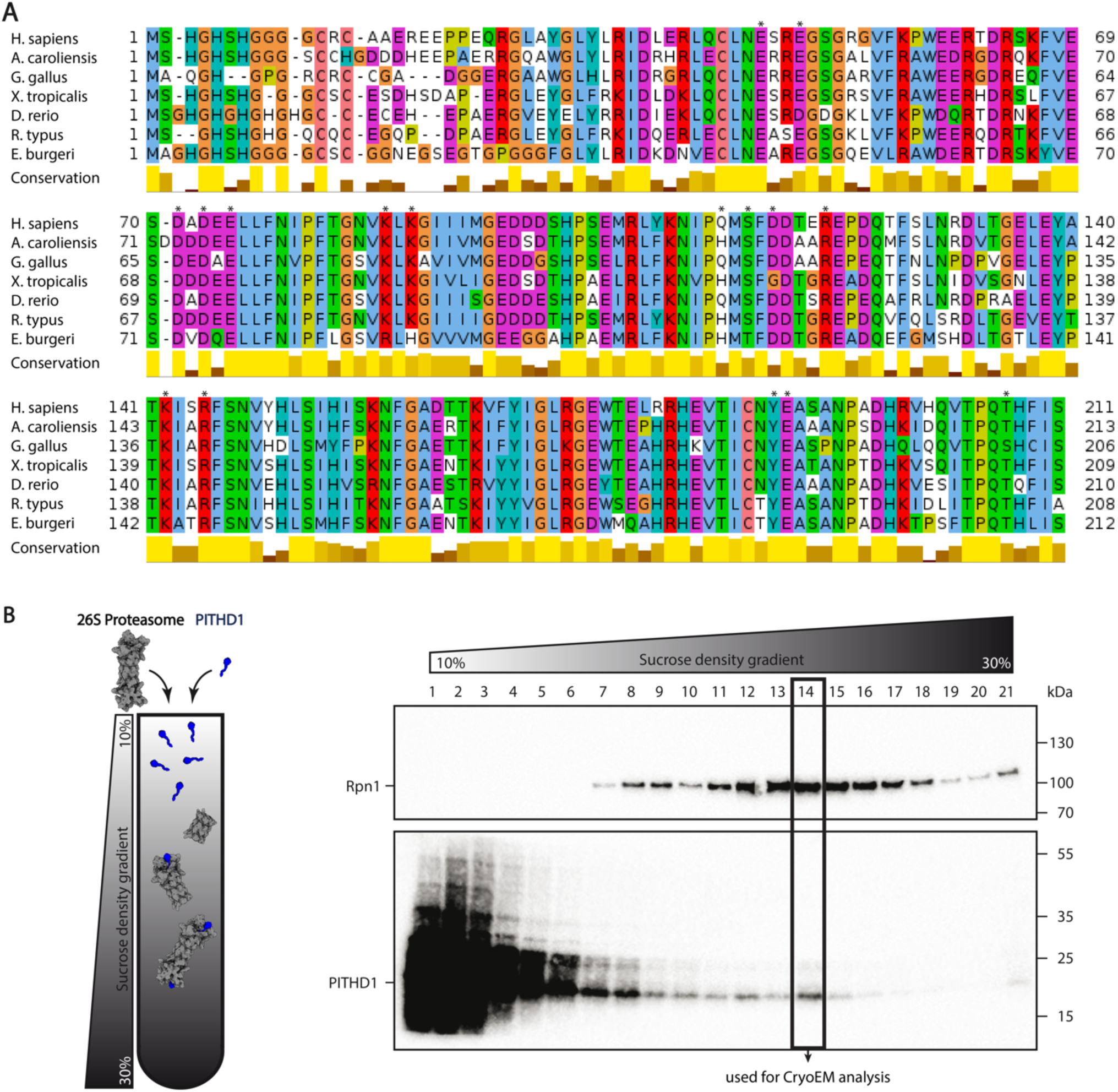
PITHD1 conservation and reconstitution. (**A**) PITHD1 conservation among Vertebrates. Asterisks indicate interactions with the 26S proteasome. (**B**) 26S proteasome - PITHD1 complex reconstitution via sucrose density gradient ultracentrifugation and WB analysis.

**Figure S3.**
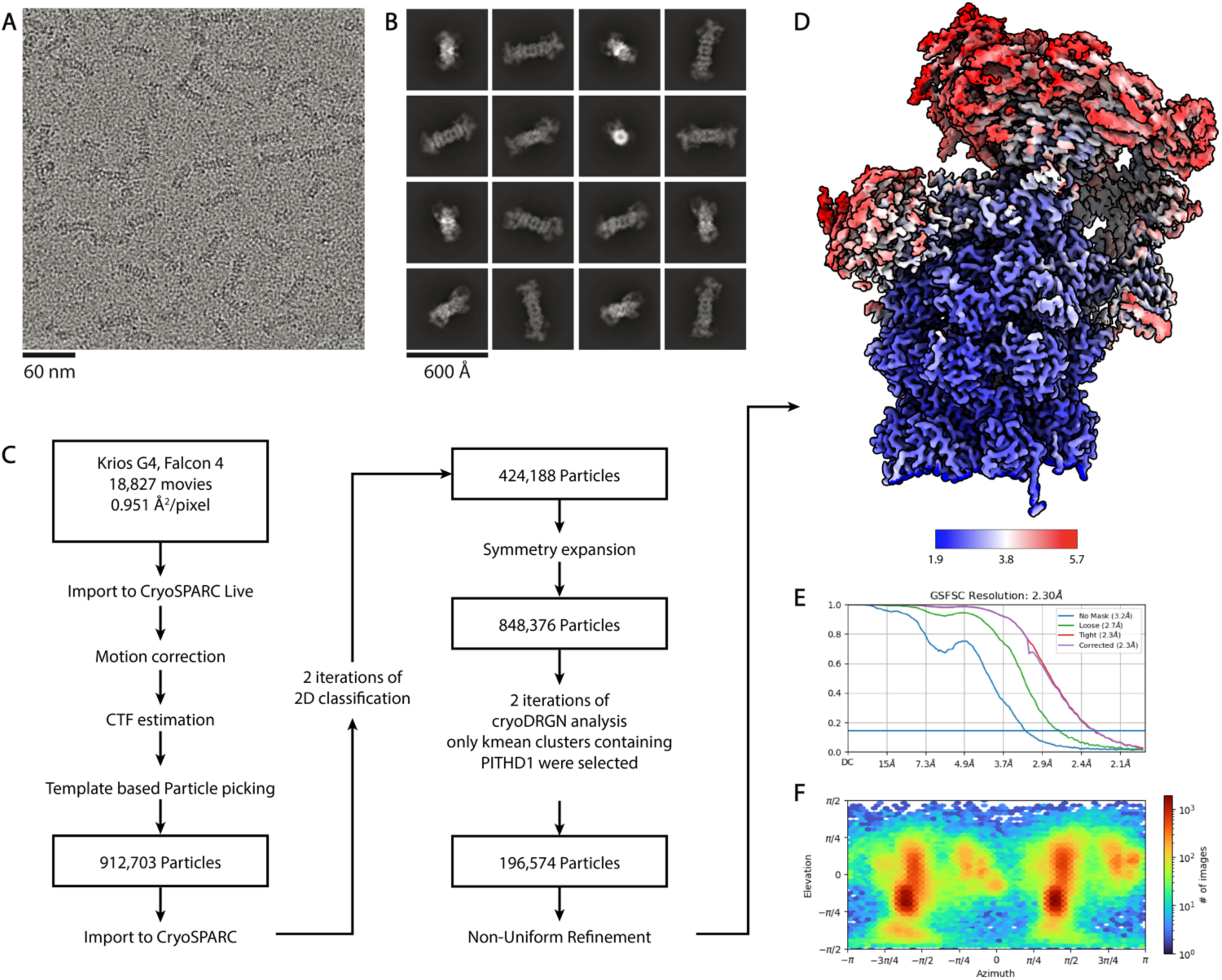
CryoEM analysis of 26S proteasome - PITHD1 complex in the S1 state. (**A**) Representative micrograph. (**B**) Representative 2D class averages. (**C**) Processing pipeline. (**D**) Local resolution map of global refined 26S proteasome - PITHD1 complex in the S1 state and corresponding (**E**) GSFSC curve and (**F**) viewing angle distribution.

**Figure S4.**
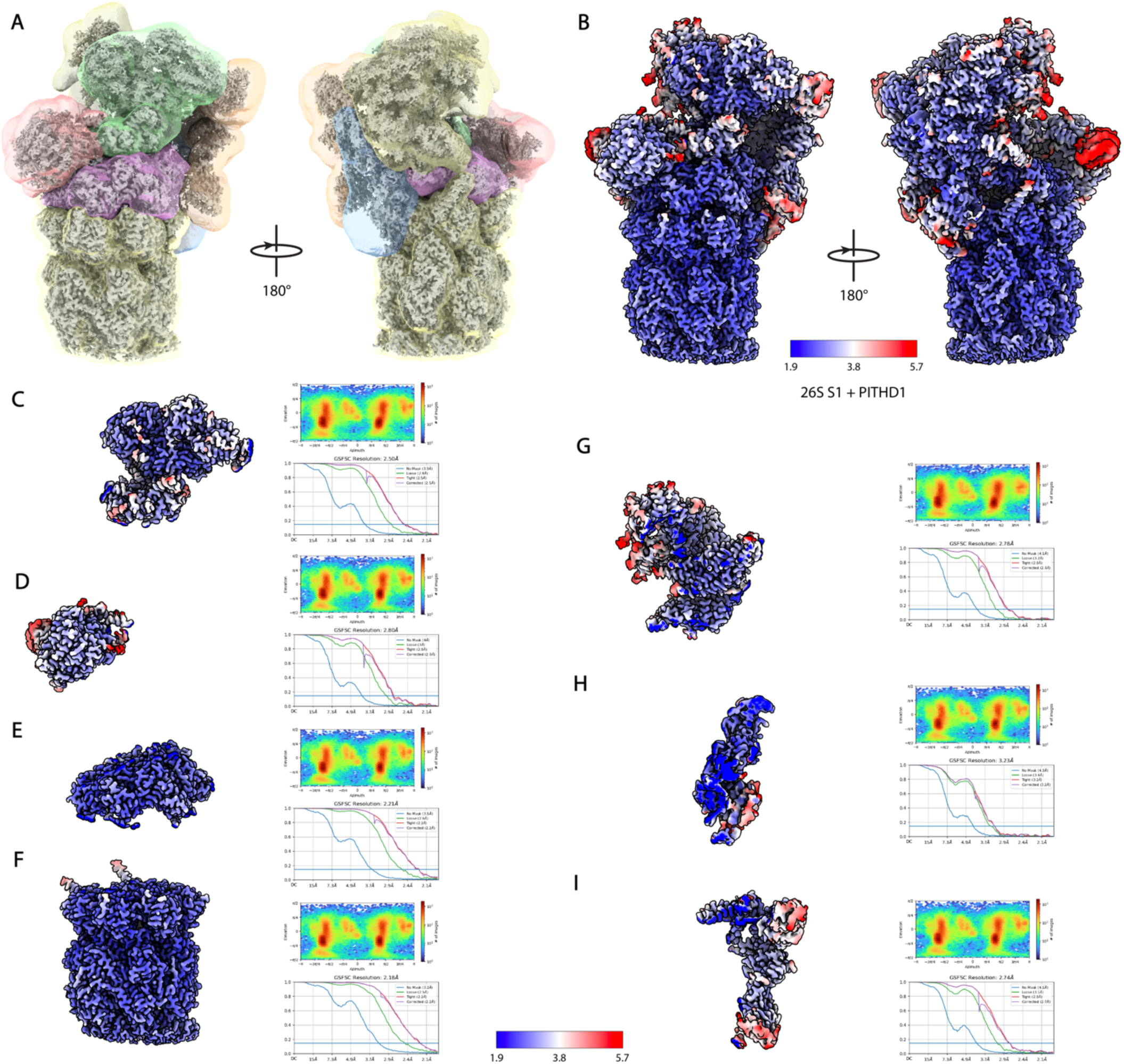
Local refinements for the 26S proteasome - PITHD1 complex in the S1 state. (**A**) Seven overlapping maps used for the local refinements. (**B**) Reconstituted local resolution maps of locally refined 26S proteasome - PITHD1 complex in the S1 state. (**C-I**) Individual local resolution estimations of local refinement maps, viewing angle distributions and GSFSC curves.

**Figure S5.**
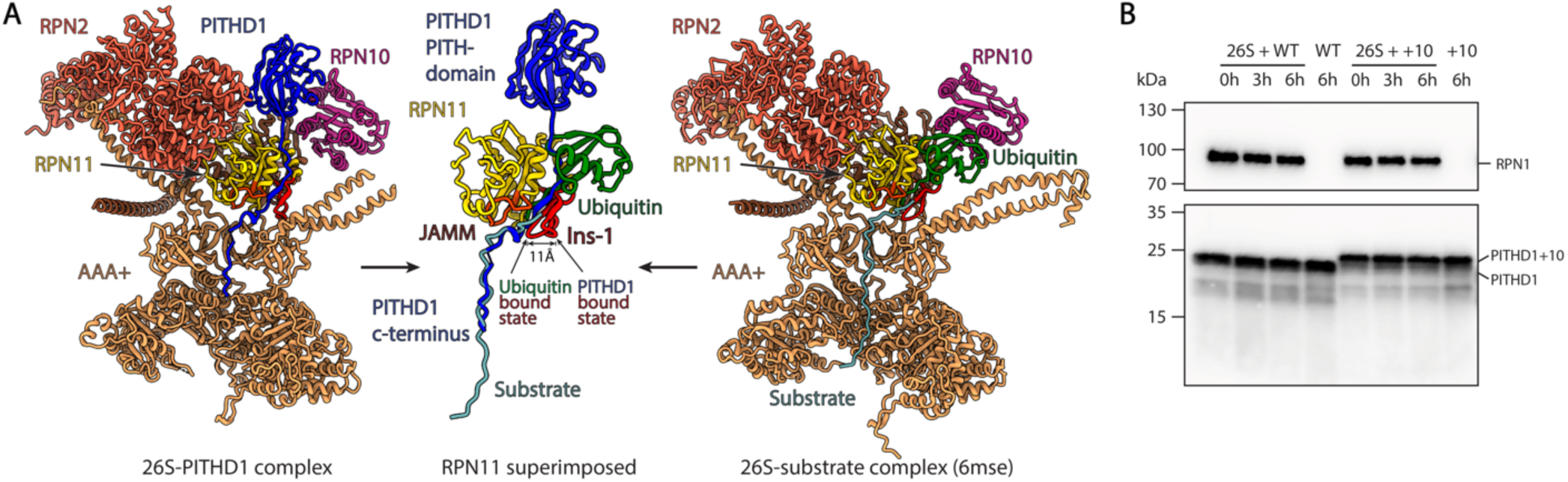
PITHD1 compared to proteasomal substrate. (**A**) 26S proteasome - PITHD1 complex compared to the substrate bound 26S proteasome in the S2 state. The binding of PITHD1 and ubiquitinated substrate is sterically exclusive. (**B**) Degradation assay with 26S proteasome and PITHD1 WT or PITHD1+10. No degradation of PITHD1 WT or PITHD1+10 can be observed.

**Figure S6.**
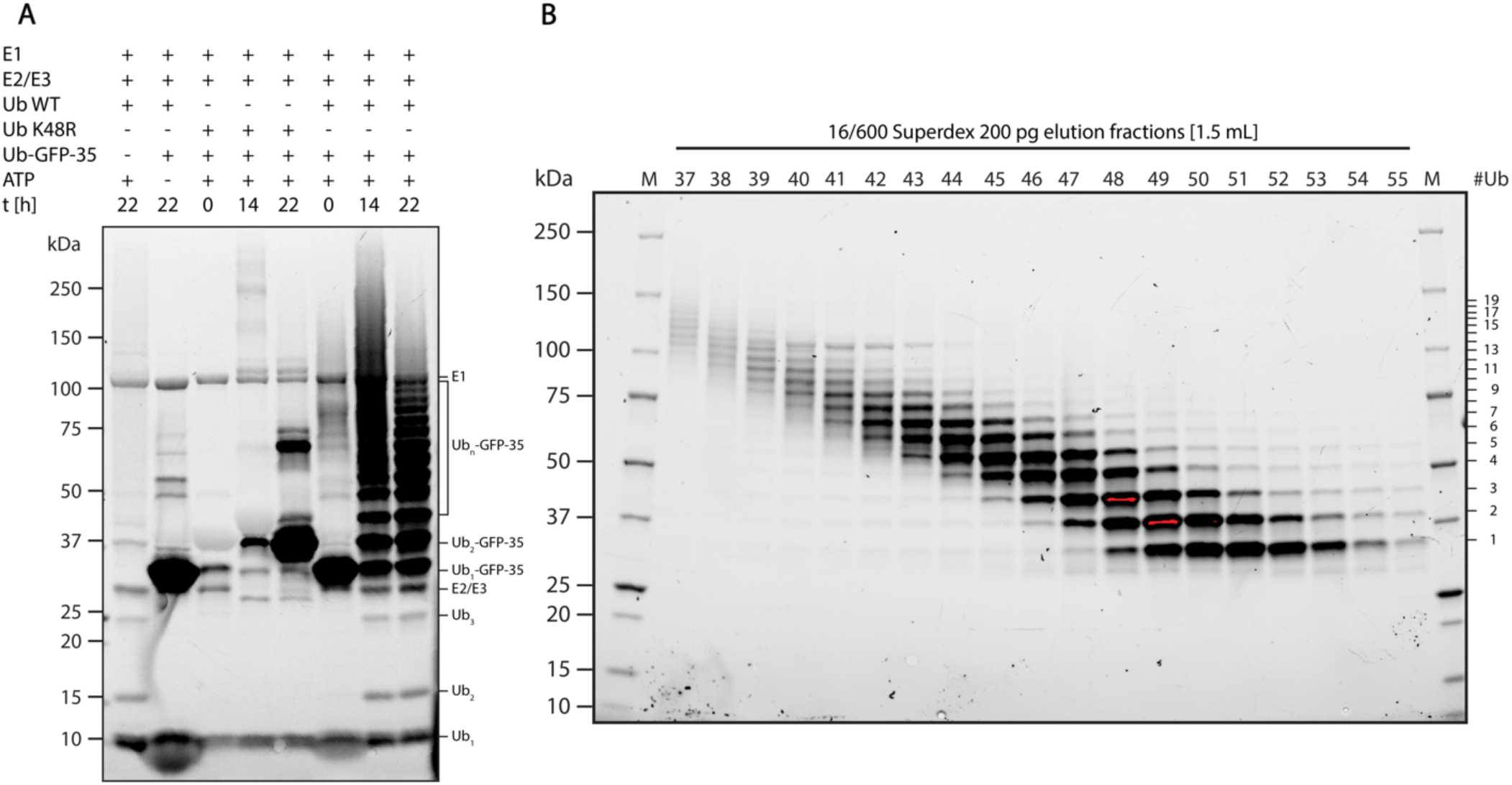
Ubiquitination of polyUb[K48]-sfGFP(CP8)-35. (**A**) SDS-PAGE of ubiquitination reaction of Ub-sfGFP(CP8)-35 with Uba1, Super E2, ubiquitin WT or ubiquitin K48R as control. A clear preference for K48 linkage can be observed. (**B**) SDS-PAGE of size exclusion chromatography of ubiquitinated Ub-sfGFP(CP8)-35 to separate the substrate according to the ubiquitin chain length. Only species with at least 4 ubiquitin moieties were used for assays or reconstitution reactions.

**Figure S7.**
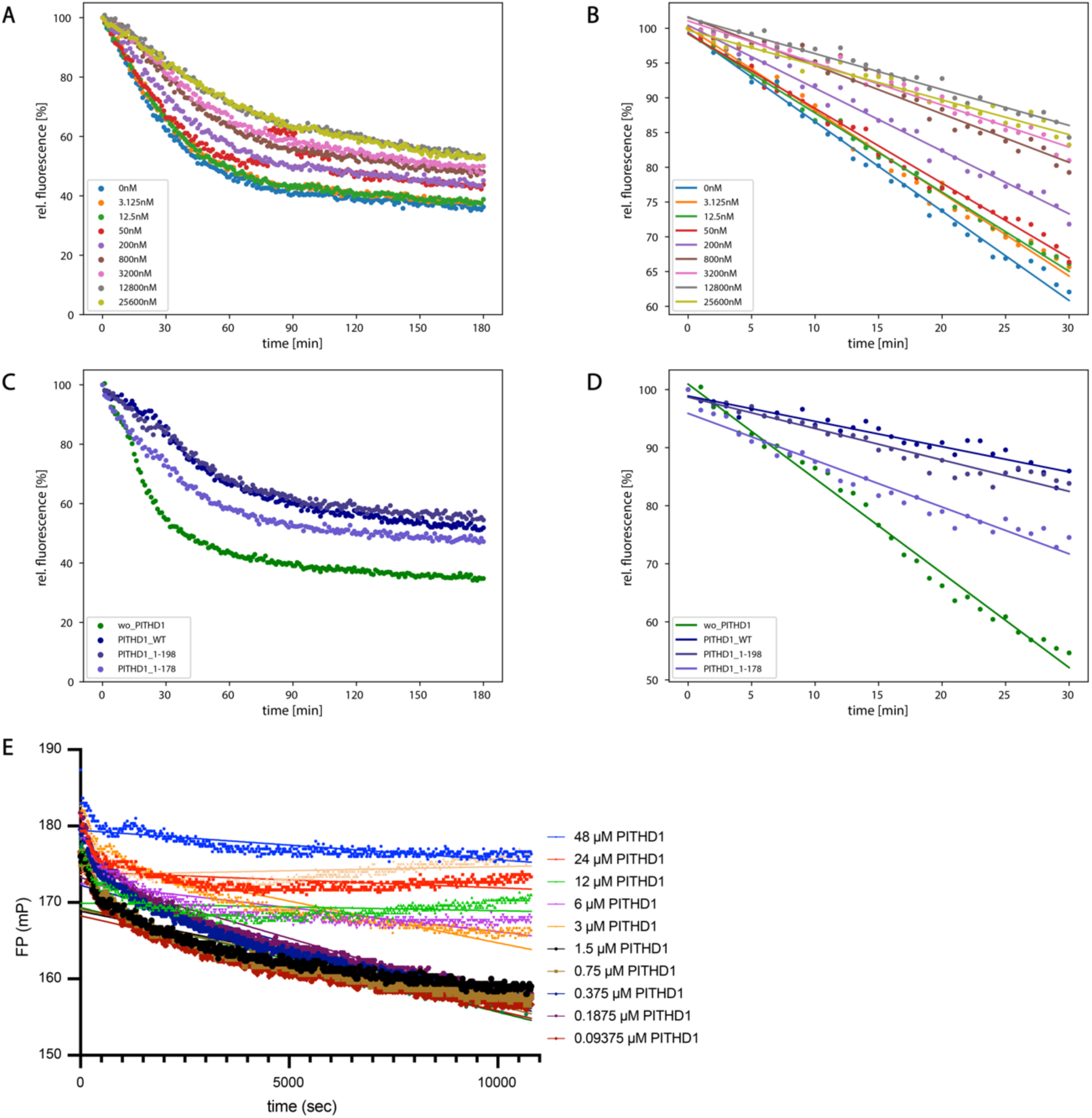
26S proteasome degradation assays. (**A**) Substrate degradation activities of 26S proteasome and different concentrations of PITHD1 WT, determined using polyUb[K48]-sfGFP(CP8)-35. (**B**) Linear fits of (**A**) to determine the initial degradation rates. (**C**) Substrate degradation activities of 26S proteasome and PITHD1 WT, PITHD1 1-198 or PITHD1 1-178, determined using polyUb[K48]-sfGFP(CP8)-35. (**D**) Linear fits of (**C**) to determine the initial degradation rates. (**E**) Deubiquitination assay of RPN11/8 heterodimer using Ub-TAMRA.

**Figure S8.**
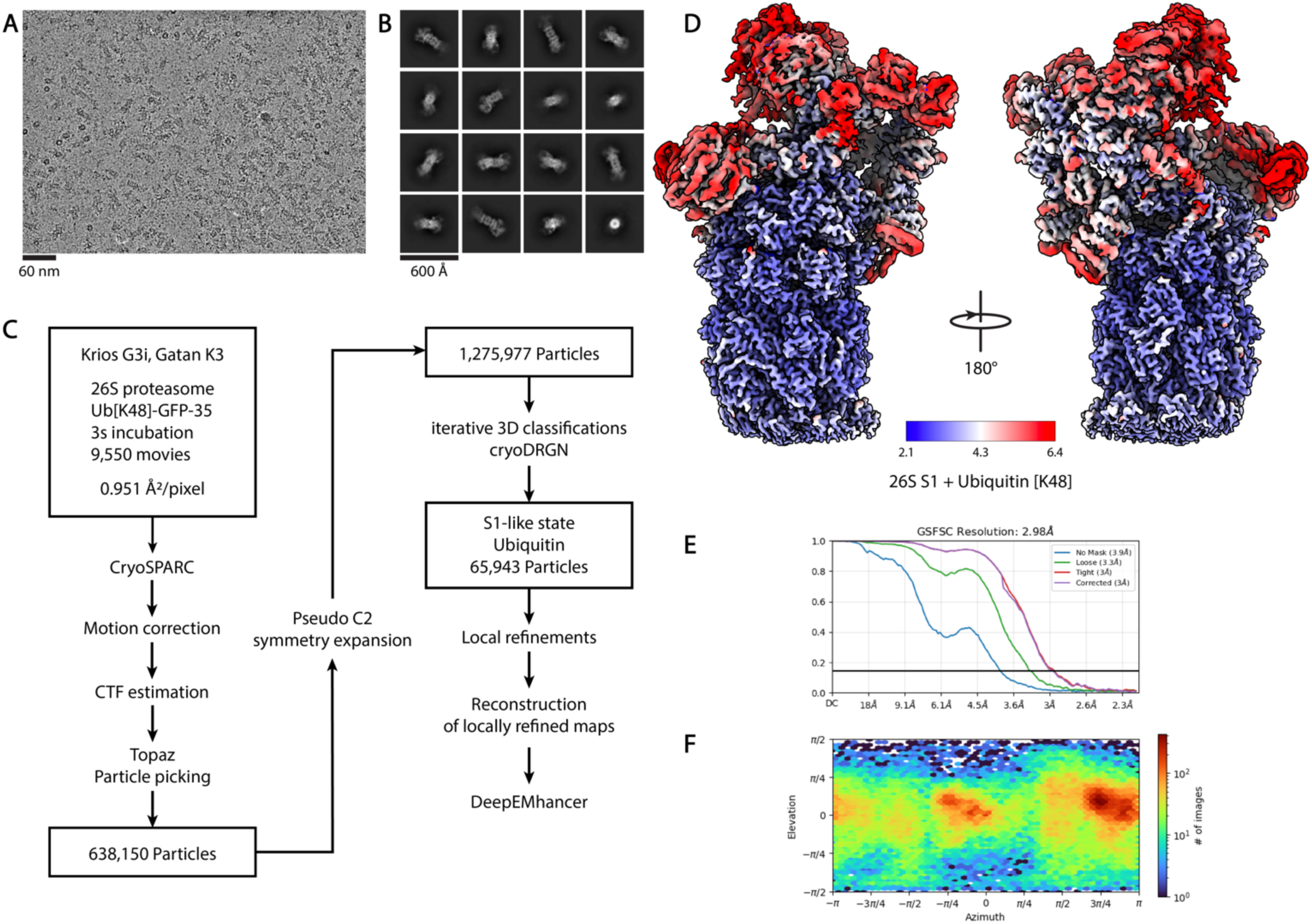
CryoEM analysis of 26S proteasome - Ub [K48] complex in the S1 state. (**A**) Representative micrograph. (**B**) Representative 2D class averages. (**C**) Processing pipeline. (**D**) Local resolution consensus map of local refined 26S proteasome - Ubiquitin [K48] complex in the S1 state and corresponding (**E**) GSFSC curve of global refined map and (**F**) viewing angle distribution.

**Figure S9.**
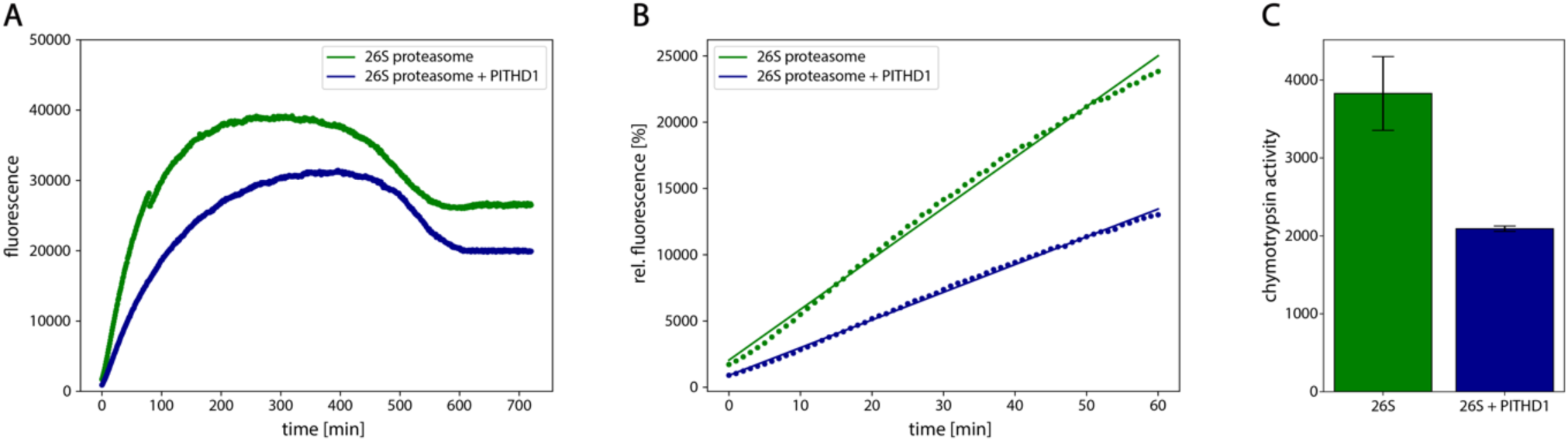
Chymotrypsin like activity of 26S proteasome dependent on PITHD1. (**A**) Chymotrypsin activities of 26S proteasome with or without added PITHD1, determined using Suc-LLVY-AMC. (**B**) Linear fits of (**A**) to determine the initial degradation rates. (**C**) Initial degradation rates of 26S proteasome with or without PITHD1.

**Figure S10.**
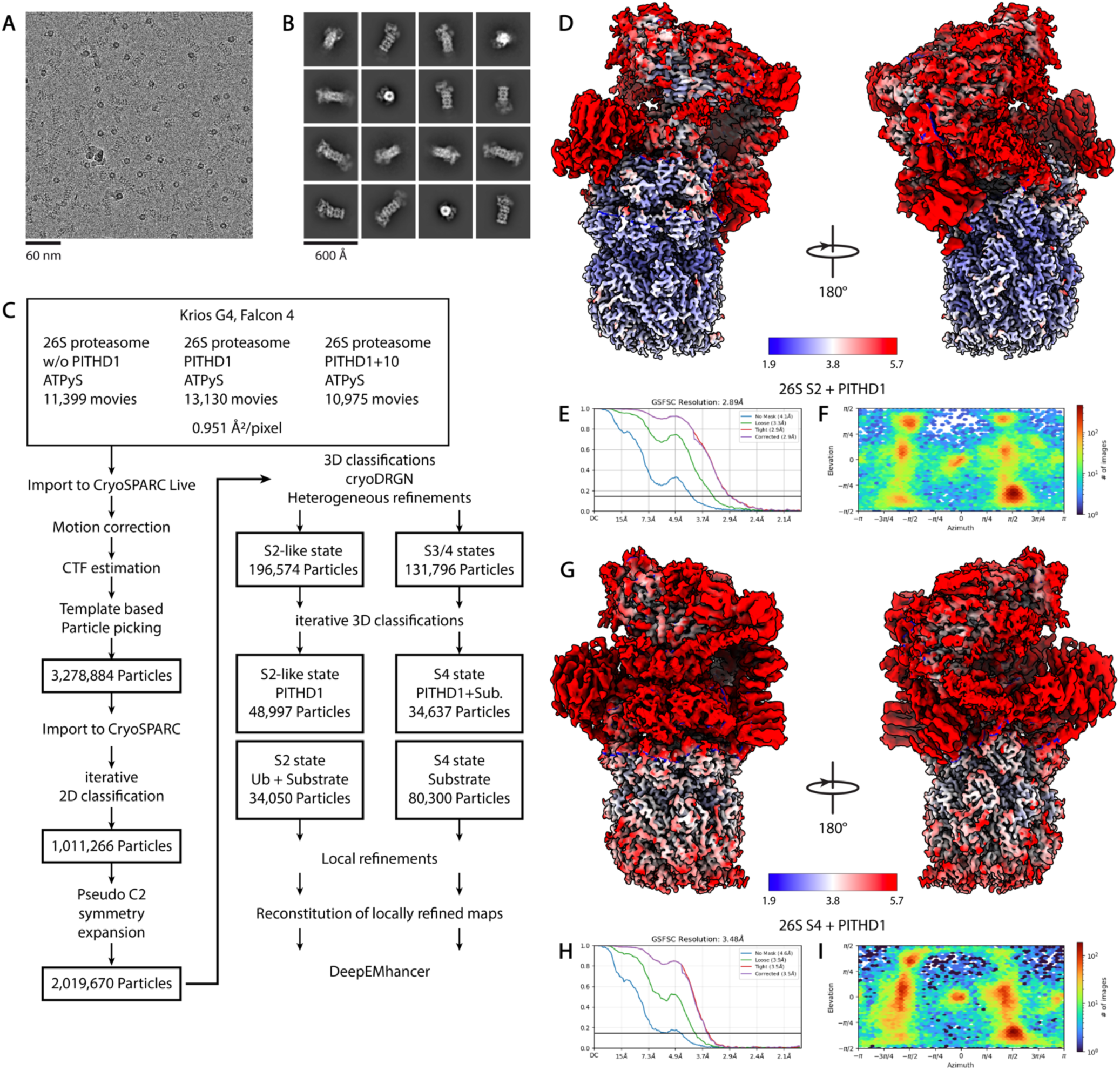
CryoEM analysis of 26S proteasome - PITHD1 complex in the S2 like and S4 states. (**A**) Representative micrograph. (**B**) Representative 2D class averages. (**C**) Processing pipeline. (**D**) Local resolution consensus map of local refined 26S proteasome - PITHD1 complex in the S2-like state and corresponding (**E**) GSFSC curve of global refined map and (**F**) viewing angle distribution. (**G**) Local resolution consensus map of local refined 26S proteasome - PITHD1 complex in the S4 state and corresponding (**H**) GSFSC curve of global refined map and (**I**) viewing angle distribution.

**Figure S11.**
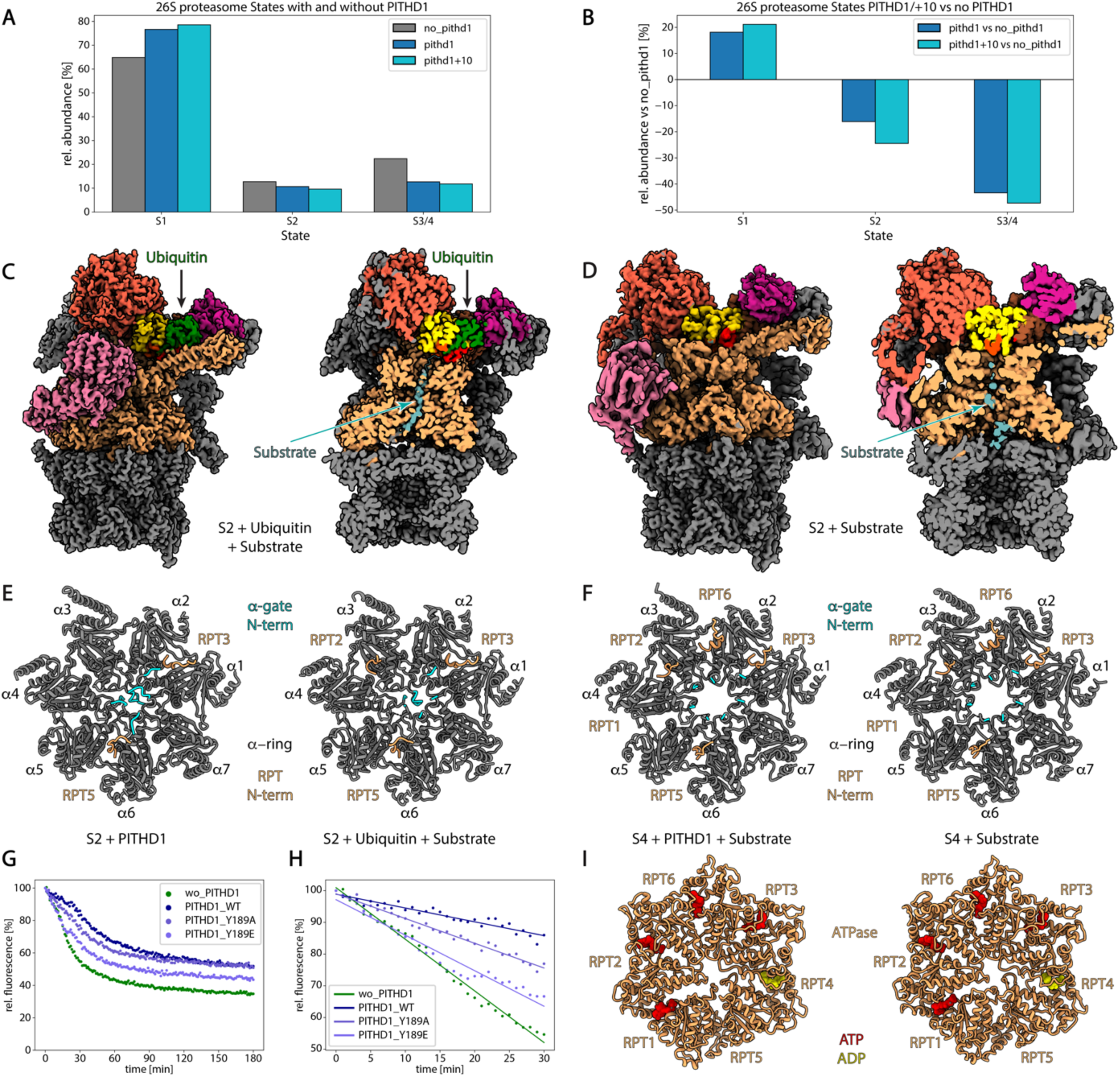
PITHD1 interactions with the activated 26S proteasome. (**A**) 26S proteasomal state distribution, depending on PITHD1 WT or PITHD1+10. (**B**) Difference plot representation of (**A**). (**C**) 26S map in the endogenous ubiquitin and substrate bound S2 state. (**D**) 26S map in the endogenous substrate bound S4 state. (**E**) HbYX insertion into α-Ring pockets and α-Ring gate states of the PITHD1 or ubiquitin and substrate bound S2 state proteasomes. (**F**) HbYX insertion into α-Ring pockets and α-Ring gate states of the PITHD1 or substrate bound s4 state proteasomes. (**G**) Substrate degradation activities of 26S proteasome and PITHD1 WT, PITHD1 Y189A or PITHD1 Y189E, determined using polyUb[K48]-sfGFP(CP8)-35. (**H**) Linear fits of (**G**) to determine the initial degradation rates. (**I**) Nucleotide states of the PITHD1 and substrate or substrate bound S state proteasomes.

---

**Table S1.**
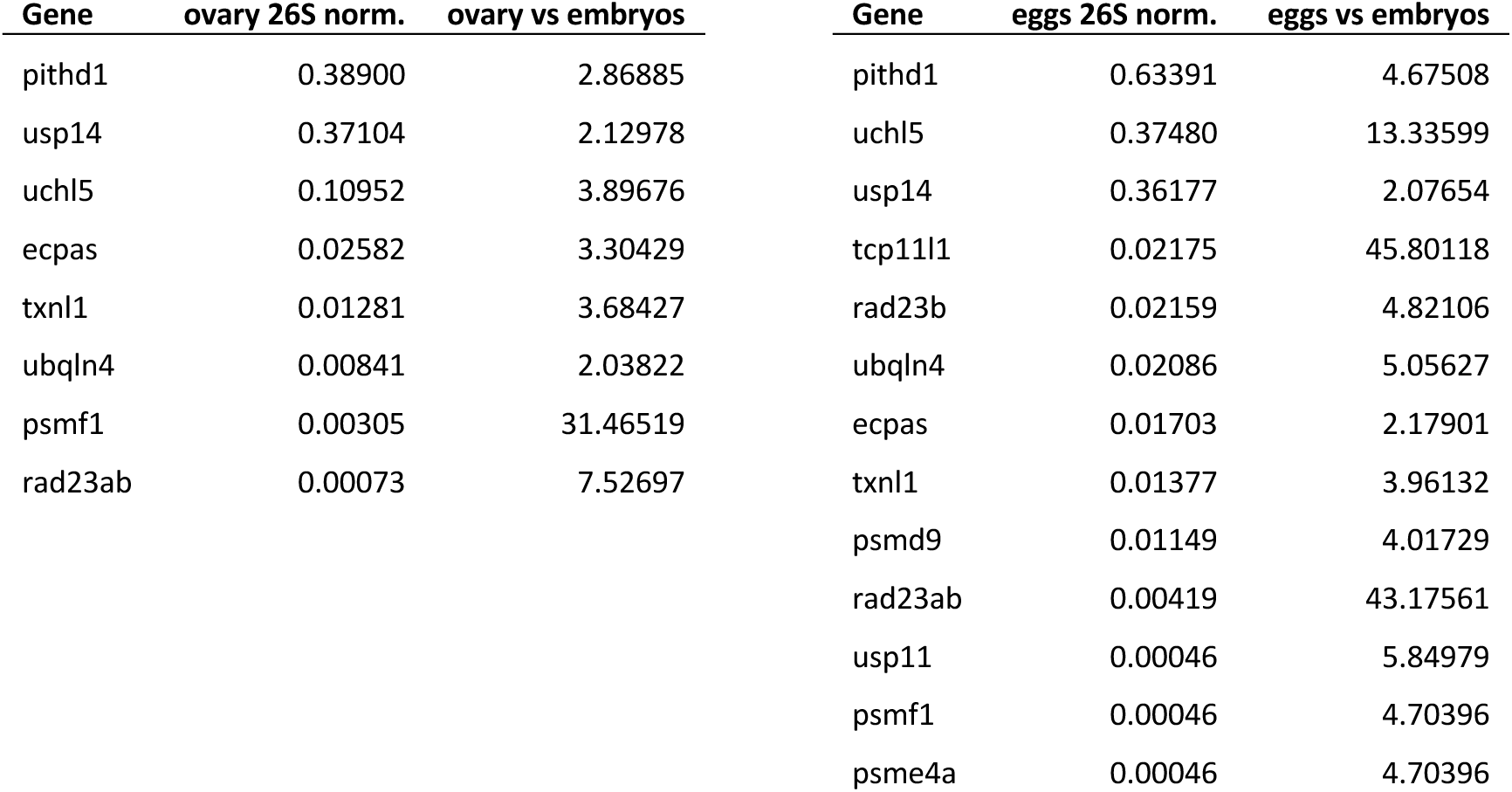
Two-fold up-regulated 26S proteasomal cofactors of ovary and eggs compared to proliferating embryos.

| Gene | ovary 26S norm. | ovary vs embryos | Gene | eggs 26S norm. | eggs vs embryos |
| --- | --- | --- | --- | --- | --- |
| pithd1 | 0.38900 | 2.86885 | pithd1 | 0.63391 | 4.67508 |
| usp14 | 0.37104 | 2.12978 | uchl5 | 0.37480 | 13.33599 |
| uchl5 | 0.10952 | 3.89676 | usp14 | 0.36177 | 2.07654 |
| ecpas | 0.02582 | 3.30429 | tcp11l1 | 0.02175 | 45.80118 |
| txnl1 | 0.01281 | 3.68427 | rad23b | 0.02159 | 4.82106 |
| ubqln4 | 0.00841 | 2.03822 | ubqln4 | 0.02086 | 5.05627 |
| psmf1 | 0.00305 | 31.46519 | ecpas | 0.01703 | 2.17901 |
| rad23ab | 0.00073 | 7.52697 | txnl1 | 0.01377 | 3.96132 |
|  |  |  | psmd9 | 0.01149 | 4.01729 |
|  |  |  | rad23ab | 0.00419 | 43.17561 |
|  |  |  | usp11 | 0.00046 | 5.84979 |
|  |  |  | psmf1 | 0.00046 | 4.70396 |
|  |  |  | psme4a | 0.00046 | 4.70396 |

**Table S2.**
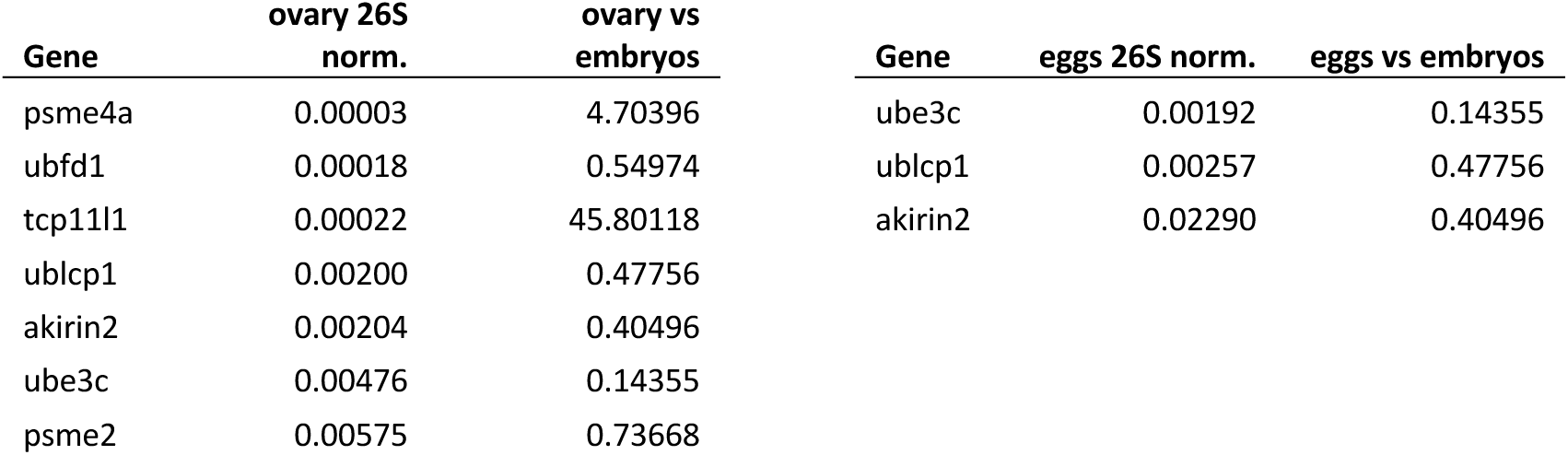
Two-fold down-regulated 26S proteasomal cofactors of ovary and eggs compared to proliferating embryos.

**Table S3.**
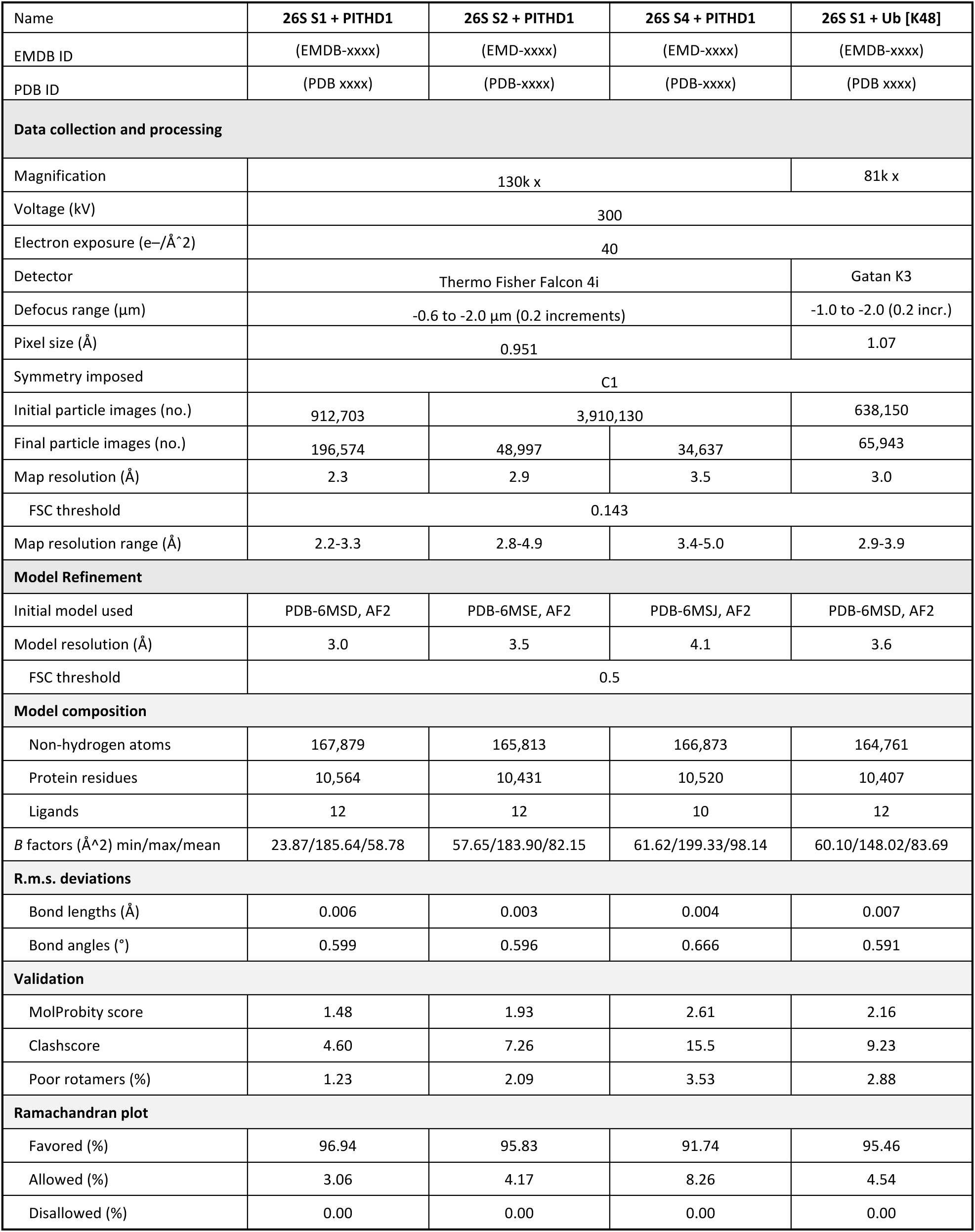
CryoEM data collection, refinement and model validation.

**Movie S1. 26S proteasomal conformational changes during substrate processing.** A morph of pdb models 6MSB, 6MSD, 6MSE, 6MSG, 6MSH illustrates the conformational changes of the 26S proteasome during substrate processing. The C-terminus of the proximal ubiquitin (green) of a substrate stabilises the Ins-1 loop (red) of RPN11 in an inward position. This facilitates the conformational change from the S1/S2 states to the active S3/S4 states by enabling the RPT4/5 coiled-coil motif (light brown) to pass through.

**Movie S2. 26S proteasomal conformational change is sterically inhibited by PITHD1.** A morph of pdb models 6MSB, 6MSD, 6MSE, 6MSG, with ubiquitinated substrate replaced by PITHD1 reveals how PITHD1’s C-terminus inhibits the conformational transition of the 26S proteasome. The C-terminus of PITHD1 (blue) stabilizes the Ins-1 loop (light blue) of RPN11 in an outward position. This restricts the conformational change from the S1/S2 states to the active S3/S4 states by causing steric clashes with the RPT4/5 coiled-coil motif (light brown).

**Movie S3. AF3 predicts a release of PITHD1’s C-terminus upon Y189 phosphorylation.** A morph derived from AF3 predictions of the functional 19S RP with ubiquitin and either PITHD1 WT or pY189 reveals that the C-terminus of PITHD1 pY189 dissociates with RPN11’s active site, thereby allowing ubiquitin to interact with this site.

